# Chronic benzodiazepine treatment triggers gephyrin scaffold destabilization and GABA_A_R subsynaptic reorganization

**DOI:** 10.1101/2025.06.02.653073

**Authors:** Caitlyn A. Chapman, Nadya Povysheva, Tyler B. Tarr, Jessica L. Nuwer, Stephen D. Meriney, Jon W. Johnson, Tija C. Jacob

## Abstract

Benzodiazepines (BZDs) are important clinical drugs with anxiolytic, anticonvulsant, and sedative effects mediated by potentiation of inhibitory GABA type A receptors (GABA_A_Rs). Tolerance limits the clinical utility of BZDs, yet the mechanisms underlying tolerance after chronic exposure have not been thoroughly investigated. Here, we assessed the impact of chronic (7-day) treatment with the BZD diazepam (DZP) on the dynamic plasticity and subsynaptic organization of the gephyrin scaffold and γ2 subunit-containing GABA_A_Rs in primary neurons. After functional confirmation of diminished BZD sensitivity, we provide the first super-resolution analysis of inhibitory nanoscale plasticity induced by chronic BZD exposure: gephyrin subsynaptic domains were smaller and the inhibitory postsynaptic area was overall diminished by DZP treatment, resulting in a condensation of synaptic γ2-GABA_A_Rs into smaller subsynaptic areas. Using a novel fluorescence-based *in situ* proximity ligation assay and biochemical fractionation analysis, the mechanism for gephyrin downregulation was revealed to be dependent on phosphorylation and protease cleavage. Accordingly, DZP treatment impaired gephyrin synaptic stability, demonstrated by live-imaging photobleaching experiments. Despite the loss of BZD sensitivity and stable synaptic gephyrin, 7-day DZP treatment did not reduce the surface or total protein levels of BZD-sensitive γ2-GABA_A_Rs, as shown in prior short-term BZD treatment studies. Instead, chronic DZP treatment induced an accumulation of γ2-GABA_A_Rs in the extrasynaptic membrane. Surprisingly, γ2-GABA_A_R interactions with gephyrin were also enriched extrasynaptically. An identified rise in extrasynaptically-localized gephyrin cleavage fragments may function to confine receptors away from the synapse, as supported by a decrease in extrasynaptic γ2-GABA_A_R mobility. Altogether, we find that chronic BZD treatment triggers several subtle converging plasticity events at inhibitory synapses which effectively restrict the synaptic renewal of BZD-sensitive GABA_A_Rs via mechanisms distinct from those observed with short-term treatment.

## Section 1: Introduction

In the central nervous system, fast inhibitory neurotransmission is primarily mediated by GABA type A receptors (GABA_A_Rs), heteropentameric chloride channels which play an important role in the maintenance and control of neuronal excitability. As neurological disorders are often characterized by an imbalance in neuronal activity, GABA_A_Rs are a key pharmacological target for widely used clinical drugs, including anesthetics, neurosteroids, barbiturates, and benzodiazepines. Benzodiazepines (BZDs) are central nervous system depressants which have persisted for decades as some of the most prescribed drugs worldwide (Kurko et al., 2015; Bachhuber et al., 2016; Maust et al., 2019). These high-efficacy, low-toxicity drugs produce anxiolytic, anticonvulsant, myorelaxant, and sedative effects through positive allosteric modulation of GABA_A_Rs and potentiation of inhibitory neurotransmission. Administration of BZDs for longer than 2-4 weeks results in tolerance to most of the behavioral effects, severely limiting clinical utility. Much of our current understanding of BZD tolerance is limited to acute or short-term BZD applications, which promote various signaling cascades that alter GABA_A_R trafficking, decrease synaptic expression, and reduce inhibitory function (Jacob et al., 2012; Nicholson et al., 2018; Lorenz-Guertin et al., 2019; González Gómez et al., 2023). Few studies have performed detailed mechanistic analysis of GABAergic signaling after long-term BZD treatment, and it remains unclear whether prolonged BZD exposure induces similar neuroplasticity. Given the persistently high patient population with long-term BZD use (Kurko et al., 2015; Olfson et al., 2015; Kaufmann et al., 2018; Tanguay Bernard et al., 2018) and high rates of patient relapse (Morin et al., 2005; Gerlach et al., 2019; Chapoutot et al., 2021), there is an urgent need to understand the impact of extended BZD treatment on inhibitory synapse plasticity and regulation.

The strength of synaptic inhibition is principally determined by GABA_A_R abundance at postsynaptic sites and receptor subunit composition, with the predominant synaptic receptor subtype consisting of two α, two β, and one γ2 subunit (Olsen and Sieghart, 2008, 2009). Dynamic trafficking mechanisms, posttranslational modifications, and regulatory protein-protein interactions further permit fine-tuning of synaptic strength (Jacob et al., 2008; Petrini and Barberis, 2014; Mele et al., 2016). GABA_A_Rs exhibit a high rate of surface lateral mobility in the plasma membrane (Choquet and Triller, 2013) but are trapped at postsynaptic sites through transient interactions with the inhibitory scaffold gephyrin, which directly binds to GABA_A_R α(1-3,5) and β(2,3) subunits via a receptor intracellular domain motif (Tretter et al., 2008, 2011; Mukherjee et al., 2011; Kowalczyk et al., 2013; Brady and Jacob, 2015; Renner et al., 2012). BZDs allosterically bind to γ2 subunit-containing GABA_A_Rs at the extracellular interface of γ2 and an α(1,2,3, or 5) subunit (Pritchett et al., 1989; Malherbe et al., 1990; Günther et al., 1995). Interestingly, acute BZD application stabilizes synaptic GABA_A_Rs in a manner dependent on gephyrin (Gouzer et al., 2014; Lévi et al., 2015), implying a conformational link between the gephyrin and BZD binding domains on GABA_A_Rs. Gephyrin is a core structural component of the inhibitory postsynaptic density critical for proper synaptic assembly and maintenance (Essrich et al., 1998; Kneussel et al., 1999; Carricaburu et al., 2024). Disruptions to gephyrin expression or synaptic stability consequently impair GABA_A_R synaptic clustering, increase GABA_A_R lateral diffusion, and impair inhibition (Jacob et al., 2005; van Zundert et al., 2005; Yu et al., 2007; Olah et al., 2023). Thus, the gephyrin-GABA_A_R interaction is essential to the regulation of inhibitory synaptic strength and, importantly, is subject to activity-dependent regulation (Petrini et al., 2014; Petrini and Barberis, 2014; Barberis, 2020; Pizzarelli et al., 2020).

Despite this central importance of gephyrin in the maintenance and plasticity of synaptic GABA_A_Rs, the impact of long-term BZD treatment on gephyrin has been severely understudied. While we and others have shown that short-term (<24 hour) BZD exposure reduces gephyrin membrane and total expression and accelerates synaptic gephyrin dynamics (Vlachos et al., 2013; Lorenz-Guertin et al., 2019), it is unknown whether these perturbations persist under conditions of more prolonged BZD treatments. In contrast to short-term treatments, we have reported similar gephyrin synaptic and total protein expression in mice after 7-day BZD treatment while extrasynaptic gephyrin levels were elevated (Lorenz-Guertin et al., 2023). Conversely, a separate investigation found decreased gephyrin mRNA levels after 7-day BZD treatment in mice, though protein expression was not assessed (Wright et al., 2014). No further studies have performed detailed analysis of chronic BZD-induced alterations to gephyrin dynamics and regulation, leaving much to be understood.

Similarly, available evidence suggests distinct mechanisms by which GABA_A_Rs are altered after long-term versus short-term BZD applications. In particular, short-term BZD treatment downregulates γ2-GABA_A_Rs and reduces miniature inhibitory postsynaptic currents (Jacob et al., 2012; Nicholson et al., 2018; Lorenz-Guertin et al., 2019), while inhibition is functionally preserved upon longer BZD exposure both *in vitro* (Hu and Ticku, 1994; Gao and Greenfield, 2005) and *in vivo* (Lorenz-Guertin et al., 2023). These findings therefore suggest that the initial adaptations occurring immediately in response to BZD application are not maintained throughout continued, long-term BZD exposure.

Here, we examined the impact of chronic DZP treatment on gephyrin and GABA_A_R nanoscale organization, regulatory processing, protein interactions, and trafficking dynamics following chronic (7-day) treatment with the BZD diazepam (DZP) in primary rodent neurons. After first confirming the development of tolerance functionally, we utilized DNA Points Accumulation in Nanoscale Topography (DNA-PAINT), a localization-based super-resolution microscopy method providing tens of nanometer spatial resolution (Jungmann et al., 2010), to provide the first analysis of the inhibitory synaptic nanostructure following chronic BZD exposure. This revealed a subsynaptic and total synapse shrinkage of gephyrin induced by chronic DZP treatment, while γ2-GABA_A_Rs were condensed into a smaller postsynaptic area. The loss of synaptic gephyrin, paralleled by a decrease in total protein expression, was associated with increased phosphorylation, protease-mediated cleavage, and reduced stability of gephyrin at synapses. This occurred alongside an enrichment of γ2-GABA_A_Rs extrasynaptically without changes to surface levels or total receptor expression. Surprisingly, these extrasynaptic γ2-GABA_A_Rs exhibited reduced mobility after chronic DZP treatment, which we show may be mediated by enhanced extrasynaptic gephyrin-GABA_A_R interactions. Altogether, we uncover multiple complementary mechanisms triggered by chronic DZP treatment that sufficiently disrupt the synaptic prevalence and renewal of BZD-sensitive GABA_A_Rs to diminish BZD potentiation of inhibition.

## Section 2: Materials and Methods

### Section 2.1: Materials, antibodies, and DNA constructs

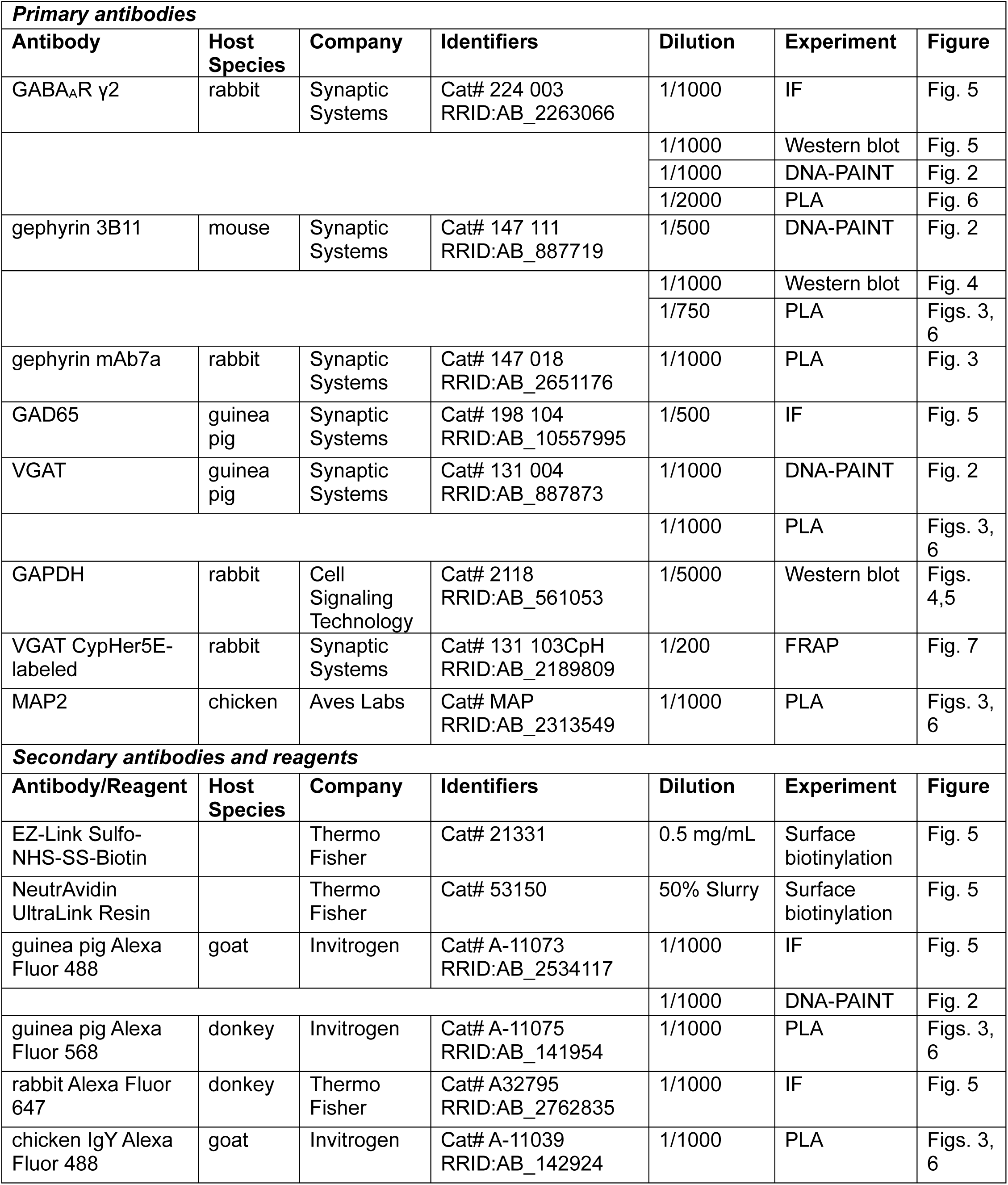
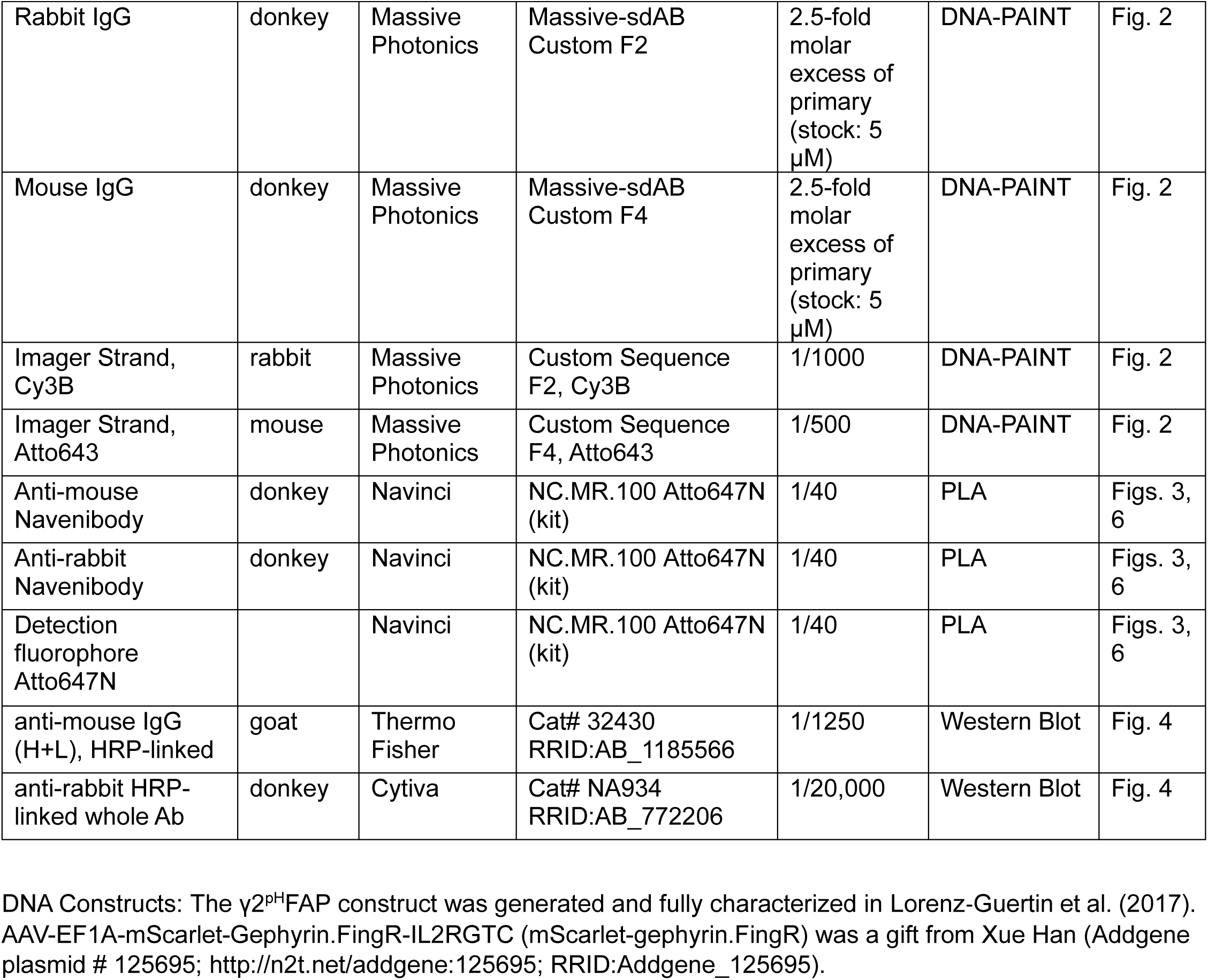

### Section 2.2: Primary neuron culture and drug treatments

All procedures were approved by the University of Pittsburgh Institutional Animal Care and Use Committee. Cortical or hippocampal neuronal cultures were prepared using procedures similar to those previously described (Jacob et al., 2005; Sahu et al., 2019). Briefly, cortical or hippocampal tissue was dissected from embryonic day 18 Sprague Dawley rats and dissociated with papain and trypsin inhibitor before resuspension in Neurobasal Media supplemented with B27 (Gibco). For FRAP experiments, neurons were nucleofected (Lonza) at plating with mScarlet-gephyrin.FingR (Gross et al., 2013; Bensussen et al., 2020) and γ2^pH^FAP (Lorenz-Guertin et al., 2017). Neurons were then cultured undisturbed until DIV 13-15, at which point they were treated with vehicle (0.1% DMSO) or 1 μM DZP (Sigma D0899) for seven days and collected for experiments at DIV 20-22.

### Section 2.3: Electrophysiology

Whole-cell patch-clamp recordings were performed on cortical neuron cultures at DIV 20-22 following 7-day treatment with vehicle or 1 μM DZP. Pyramidal neurons were visualized by IR-DIC video microscopy and identified by their apical dendrites and large triangular soma. Patch electrodes (5–10 MΩ open-tip resistance) were filled with an intracellular solution containing (in mM): 140 CsCl, 2 MgCl_2_, 0.1 CaCl_2_, 10 HEPES, 10 phosphocreatine, 4 ATP-Mg, 0.3 GTP, and 1.1 EGTA; pH 7.25. Extracellular Ringer solution of the following composition was used (in mM): 126 NaCl, 24 NaHCO_3_, 2.5 KCl, 1.25 NaH_2_PO_4_, 1 MgSO_4_, 2 CaCl_2_, 10-20 glucose; bubbled with a 95% O_2_/5% CO_2_ gas mixture; pH ∼7.3. Current recordings were performed with a Multi-Clamp 700A amplifier (Axon Instruments, Union City, CA, USA). Signals were filtered at 2 kHz and acquired at a sampling rate of 10 kHz using Clampex 10.2 software (Molecular Devices Corporation, Sunnyvale, CA, USA). Access resistance was 10–20 MΩ and remained relatively stable during experiments (≤30% increase). Recordings were corrected for the liquid junction potential. All currents were recorded at a holding potential of −70 mV. Miniature inhibitory postsynaptic currents (mIPSCs) were recorded in the presence of NBQX (20 μM), D-APV (50 μM), and TTX (1 μM) to inhibit AMPAR, NMDAR, and voltage-gated sodium channels, respectively. Miniature events were analyzed using the MiniAnalysis Program (Synaptosoft, Decatur, GA, USA) as previously described (Povysheva and Johnson, 2016). The percent DZP potentiation of mIPSC amplitude and tau was determined by the percent change from baseline upon acute application of 1 µM DZP.

### Section 2.4: DNA-PAINT Immunostaining, Imaging, and Analysis

DIV 20-22 neurons were collected at the end of the 7-day drug treatment, rapidly washed with DPBS, and fixed for 10 minutes in PBS containing 4% paraformaldehyde (PFA) and 4% sucrose. For DNA-PAINT staining, primary antibodies against the γ2-GABA_A_R subunit and gephyrin were each separately pre-incubated for 20 minutes with custom-made single-domain secondary nanobodies coupled to oligonucleotides (Massive Photonics) such that the nanobodies were in 2.5 molar excess of the respective primary antibody (Sograte-Idrissi et al., 2020). After blocking in blocking solution (DPBS containing 10% horse serum and 0.5% bovine serum albumin (BSA)), neurons were incubated overnight with the γ2 antibody/nanobody pre-mix to identify surface γ2-GABA_A_Rs, then permeabilized for 10 minutes with blocking solution containing 0.2% Triton X-100. Following permeabilization, neurons were incubated overnight with the gephyrin antibody/nanobody pre-mix and a primary antibody against the vesicular GABA transporter VGAT, which was used to confirm synaptic localizations. Secondary antibody was added for one hour at room temperature the next day followed by a 10-minute post-fix. Dishes were stored in PBS at 4°C for up to two weeks prior to image collection.

Single molecule localization imaging was performed on an Olympus inverted microscope using a 100x TIRF oil-immersion objective (1.5 NA). The microscope was equipped with a super-resolution Abbelight 360 SAFe dual-camera (Hamamatsu Fusion) system. The incident angle was manually adjusted for Highly Inclined and Laminated Optical (HILO) illumination to achieve brightest blinking signals. Built-in TrueFocus Red Z drift was used to maintain stability in the z-focal plane throughout image collection. Imager strands diluted to their final concentration (∼1-2 nM) in PBS supplemented with 500 mM NaCl were added to prepared neuron dishes. Prior to imaging, a snapshot was taken with the 488 nm laser to identify VGAT clusters. 30,000 frames were then collected at 100 ms exposure with excitation using the 561 and 640 nm lasers. On the same day, separate dishes coated with TetraSpeck beads (Invitrogen) were imaged for 100 frames at 100 ms exposure to facilitate channel alignment.

Single molecule processing and analysis was performed using procedures similar to those previously described (Schnitzbauer et al., 2017; Chen et al., 2020). Image files were converted from .tif to .raw format in FIJI using the plugin raw-yaml exporter (https://github.com/jungmannlab/imagej-raw-yaml-export) to allow further processing in Picasso (https://github.com/jungmannlab/picasso). Picasso: Localize and Picasso: Filter were used to identify and refine localizations for each channel. Drift correction was then performed in Picasso: Render, where localizations persisting for more than one frame were linked. For each neuron, exported localizations from the two channels were then combined in Excel to facilitate import into MATLAB. Synapses were manually selected based on colocalization with VGAT, significant overlap of γ2-GABA_A_R and gephyrin, high local protein density, and a size of approximately 100 – 800 nm. Selected synapses were filtered using the MATLAB function DBSCAN according to the following parameters to remove background localizations outside of the synapse boundary: γ2-GABA_A_R epsilon=40 nm, minimum points=5; gephyrin epsilon=30 nm, minimum points=5. Areas of high local protein density (subsynaptic domains, SSDs) were then analyzed in MATLAB as previously described (Chen et al., 2020; Anderson et al., 2023). Briefly, identification of SSDs was based on having a local protein density greater than a specified threshold determined by comparison to a randomized cluster with bounding areas created using the alphaShape function (alpha radius: 150 nm). Enrichment index was defined as the average local density of protein *a* within a 60 nm range from an SSD peak of protein *b*, as previously described (Chen et al., 2020; Dharmasri et al., 2024).

### Section 2.5: Fixed Immunofluorescence (IF)

Following 7-day treatment with vehicle or 1 μM DZP, DIV 20-22 neurons were rapidly washed with DPBS then immediately fixed for ten minutes in PBS containing 4% PFA and 4% sucrose. For surface staining of γ2-GABA_A_Rs, neurons were blocked for 30 minutes then incubated under nonpermeabilized conditions with primary antibodies overnight at 4°C. Permeabilization was performed after washing the next day by 10-minute incubation with blocking solution containing 0.2% Triton X-100. This was followed by overnight intracellular staining for GAD65 at 4°C. Neuron coverslips were washed the next day, then incubated with secondary antibodies for one hour at room temperature before mounting.

### Section 2.6: Proximity Ligation Assay (PLA)

For *in situ* PLA experiments, 7-day vehicle- or DZP-treated neurons were collected at DIV 21 by rapid washing in DPBS followed by immediate fixation in PBS containing 4% PFA and 4% sucrose for 10 minutes. Neurons were then permeabilized for 10 minutes in 0.2% Triton X-100. PLA was performed according to the manufacturer’s protocol using the NaveniFlex Cell MR Atto647N kit (Navinci Diagnostics, Sweden). In brief, coverslips were blocked in kit-supplied blocking solution for one hour in a humidity chamber at 37°C then incubated with primary antibodies overnight at 4°C. Oligonucleotide-conjugated secondary antibodies (Navenibodies) were added the next day for one hour in a humidity chamber at 37°C, followed by washing and incubation in a ligase solution to permit hybridization of proximal Navenibodies. Subsequent addition of a polymerase solution containing fluorescently-labeled oligonucleotides promoted rolling circle amplification.

Next, overnight counterstaining was performed with primary antibodies against microtubule-associated protein 2 (MAP2), to facilitate visualization of neuronal dendrites, and the inhibitory presynaptic marker VGAT to identify synaptic signals. This was followed by secondary antibody incubation and DAPI nuclear staining.

### Section 2.7: IF and PLA Imaging and Analysis

Fixed images were acquired using a Nikon A1 Confocal microscope equipped with a 60x oil-immersion objective (NA 1.49) at a zoom of 2x with sequential laser scanning. Image acquisition and laser settings were kept consistent within each culture and between treatment groups with the researcher blinded to the experimental conditions before data collection and throughout data analysis. Data were analyzed using NIS Elements AR 5.30.05 Software (Nikon, NY) with binary thresholding. For immunofluorescence (IF) experiments, synaptic and extrasynaptic receptor quantification was performed as previously described (Nuwer ^e^t al., 2023^)^. Briefly, synaptic γ2-GABA_A_R signal was determined by binary intersection of the surface γ2-GABA_A_R and GAD65 thresholds, while extrasynaptic γ2-GABA_A_Rs were defined by subtraction of the synaptic γ2-GABA_A_R threshold from the surface receptor threshold. Prior to subtraction, the synaptic receptor binary threshold was dilated once. For each neuron, three 10 μm dendritic regions of interest (ROIs) were collected to analyze each binary threshold, with measurements of the number of clusters, binary area, mean intensity, and sum intensity exported for further analysis. The values of the three ROIs per cell were averaged prior to compiling. For PLA analysis, bright circular proximity ligation (PL) signals having a typical diameter of 0.50 μm, in agreement with the manufacturer-defined size of typical PL signals, were identified using Bright Spot Detection. Manual exclusion was used sparingly to remove nonspecific signals that were not localized to any visible cell processes. Synaptic PL signals were defined using the binary operation “Having,” which isolated PL spot signals containing any pixels overlapping with the VGAT threshold. Whole field and synaptic measurements were exported for further analysis. The number of extrasynaptic PL signals was computed manually in Excel by subtraction of the number of synaptic PL signals from the total (whole-field) number of PL signals. Fluorescence intensity values for IF experiments, or PL signal measurements for PLA experiments, were normalized to the vehicle average for each independent culture.

### Section 2.8: Surface biotinylation and western blotting

Surface biotinylation experiments were performed as previously described (Nuwer et al., 2021). Briefly, 7-day vehicle- or DZP-treated neurons were rapidly washed twice with DPBS supplemented with 1 mM CaCl_2_ and 0.5 mM MgCl_2_. Dishes were then incubated with 0.5 mg/mL of cell-impermeant EZ-Link Sulfo-NHS-SS-Biotin (Thermo Fisher) for 15 min at 4°C. Excess biotin was quenched by three washes with 100 mM glycine followed by one wash in DPBS. Neurons were then lysed in RIPA containing 50 mM Tris-HCl at pH 8.0, 150 mM NaCl, 1% Igepal, 0.5% sodium deoxycholate, 0.1% SDS, 1 mM EDTA, 2 mM sodium orthovanadate, 10 mM NaF, and protease inhibitor cocktail (Sigma P8340). Lysates were sonicated, solubilized for 15 min at 4°C, then centrifuged (13,000 rpm, 15 min, 4°C) to remove cell debris. After quantifying protein concentrations by BCA Protein Assay (Thermo Fisher), equal amounts of protein were incubated with NeutrAvidin UltraLink Resin (Thermo Fisher) for 90 min at 4°C with rotation. This was followed by three washes with RIPA supplemented with 500 mM NaCl and elution of isolated biotinylated surface proteins with SDS loading buffer and heating (55°C, 10 min). Surface and total protein fractions were resolved by SDS-PAGE then transferred overnight to supported nitrocellulose membrane (Bio-Rad). Membranes were incubated with primary antibodies overnight at 4°C. After washing with TBS supplemented with 1% Tween 20 (TBST) the next day, HRP-coupled secondary antibodies were added for 1 hour at room temperature followed by chemiluminescent visualization. Analysis was performed in Image Lab 6.0 (Bio-Rad) using the volume tool to quantify immunoreactivities with global background subtraction. Within each independent culture, measurements were normalized to the vehicle-treated average. The absence of GAPDH signal in the surface fraction was used to confirm surface-specific labeling.

### Section 2.9: Subcellular Fractionation and western blotting

Fractionation experiments were performed as previously described (Goebel-Goody et al., 2009; Lorenz-Guertin et al., 2023). Neurons were treated with vehicle or DZP for 7 days (∼DIV 14-21) then lysed in sucrose buffer containing (in mM): 320 sucrose, 10 Tris-HCl, 1 EDTA, 2 Na_3_VO_4_, 10 NaF, and protease inhibitor cocktail (Sigma P8340). An initial slow-speed centrifugation (1,000 x*g*, 10 min) was performed to remove nuclear debris, and a small amount of supernatant (S1) representing the total fraction was set aside for downstream analysis. Subsequent centrifugation of S1 (15,000 x*g*, 30 min) generated a cytosolic fraction (supernatant S2) and crude membrane fraction (pellet P1). P1 was resuspended in 496 μL of H_2_O containing phosphatase and protease inhibitors and incubated on ice for 15 min followed by addition of 3.75 μL of 1 M HEPES solution and another 15 min incubation. Samples were then spun at high speed (25,000 rpm, 20 min; Beckman Coulter Optima Max-E Ultracentrifuge) and the supernatant was discarded. The pellet (P2) was resuspended in sucrose buffer containing Triton X-100 (final concentration, 0.5%) and spun for 60 min at 53,000 rpm. The resulting Triton-insoluble pellet (P3), defined as the synaptic fraction, was resuspended in sucrose buffer and sonicated. SDS was then added (final concentration, 1%) to facilitate protein solubilization. The Triton-soluble supernatant (S3), defined as the extrasynaptic fraction, was concentrated by overnight incubation with 4x volumes of acetone at −20°C. The resulting precipitate was isolated by centrifugation (15,000 x*g*, 10 min), resuspended in sucrose buffer, sonicated, and SDS added to a final concentration of 1% to solubilize proteins. Fractions were frozen at −80°C until downstream analysis. All steps were performed on ice, and all centrifugations were at 4°C. Pellets were rinsed twice between centrifugation steps with sucrose buffer containing inhibitors to minimize potential contamination between fractions. Protein concentrations for each fraction were determined by BCA Protein Assay (Thermo Fisher). Equal amounts of protein were resolved by SDS-PAGE and transferred overnight to supported nitrocellulose membrane (Bio-Rad). Membranes were then processed and analyzed as in Section 2.8.

### Section 2.10: Fluorescence Recovery after Photobleaching (FRAP) Imaging and Analysis

Neurons expressing γ2^pH^FAP and mScarlet-Gephyrin.FingR were treated with vehicle or 1 μM DZP for 7 days then subjected to live-cell FRAP studies. Hippocampal neurons were used due to their improved longevity over cortical neurons following transfection. For live imaging, neurons were rapidly washed with, then transferred to, Hepes-buffered saline (HBS) imaging solution containing (in mM): 135 NaCl, 4.7 KCl, 10 Hepes, 11 glucose, 1.2 MgCl_2_, and 2.5 CaCl_2_ (adjusted to pH 7.4 with 1 N NaOH). To confirm synaptic localization of mScarlet-gephyrin.FingR clusters, live neurons were first incubated with CypHer5E-labeled VGAT for 1-2 hr to allow uptake into recycling vesicles. Experiments were performed using a Nikon A1 Confocal microscope with a 60x oil-immersion objective (NA 1.49) at 2x zoom. Stage and objective heaters were set to 37°C throughout the imaging period. Following an initial acquisition phase, 4-6 synaptic regions and 1 extrasynaptic region per neuron were subjected to photobleaching for one minute using the 488 and 561 lasers at 25% power. 10 nM MG-βTau was added to the dish immediately after photobleaching to re-identify surface γ2^pH^FAP clusters as previously described (Lorenz-Guertin et al., 2019). Images were then taken every two minutes for the next 30 minutes to monitor fluorescence recovery. γ2^pH^FAP signal was considered synaptic by colocalization with bright clusters of mScarlet-Gephyrin.FingR. Time series alignment was performed before analysis to correct for drift during image collection. Fluorescence recovery was calculated as previously described (Jacob et al., 2005) according to the following equation: (F_t_ – F_0_)/(F_i_ – F_0_), where F_0_ is the fluorescence intensity within each ROI immediately after photobleaching, F_i_ is the average fluorescence intensity prior to photobleaching, and F_t_ is the measured fluorescence at each time point following bleaching.

### Section 2.11: Statistical Analysis

Statistical analysis and graphical representation of data were performed using GraphPad Prism 10.3.1. Data were assessed for normality using D’Agostino & Pearson, Anderson-Darling, Shapiro-Wilk, and Kolmogorov-Smirnov tests. For data that passed the normality tests, two-tailed unpaired *t*-tests were performed to compare vehicle-versus DZP-treated groups; otherwise, two-tailed Mann-Whitney tests were conducted. Outliers were identified using Grubbs’ (α=0.05) or ROUT (Q=1%) and removed as appropriate. All data are presented as mean ± standard error of the mean (SEM) unless otherwise stated. Additional information on specific statistical analyses can be found in the respective figure legends or Supplementary Table S1.

## Section 3: Results

### Section 3.1: Primary cortical neurons are resistant to benzodiazepine potentiation after chronic 7-day DZP treatment

We first established a cultured neuron model of tolerance to evaluate the impact of chronic BZD treatment on basal inhibition and BZD potentiation. BZD binding in the presence of GABA stabilizes the pre-activation receptor conformation and increases the frequency of channel opening (Gielen et al., 2012; Mozrzymas et al., ^2^007^)^, thus enhancing current amplitude and prolonging inhibitory currents (higher tau of decay, τ_decay_).

Following a 7-day treatment with either vehicle (0.1% DMSO) or 1 μM DZP, whole-cell recordings were performed in primary cortical neurons to measure miniature inhibitory postsynaptic currents (mIPSCs). In agreement with our prior report (Lorenz-Guertin et al., 2023), mIPSC parameters were unaltered after long-term BZD treatment (**Fig. S1**), indicating preservation of inhibitory synapse function. BZD potentiation was then assessed by acutely applying 1 μM diazepam to 7-day vehicle- and DZP-treated neurons (**Fig. 1A**) and quantifying the corresponding potentiation of mIPSC amplitude (**Fig. 1B,C**) and τ_decay_ (**Fig. 1D,E**). As expected, acute diazepam application to 7-day vehicle-treated neurons produced a 50% increase in mIPSC amplitude from baseline (**Fig. 1C**) and a 75% percent increase in τ_decay_ (**Fig. 1E**). In contrast, 7-day DZP-treated neurons showed a substantial diminishment in acute diazepam potentiation of mIPSCs (**Fig. 1B,D**). DZP potentiation of mIPSC amplitude was nearly completely lost, reduced to only ∼7% (**Fig. 1C**), and potentiation of τ_decay_ was reduced to ∼25% (**Fig. 1E**). Thus, these results are consistent with impaired BZD sensitivity without loss of basal synaptic inhibition.

**Figure 1.**
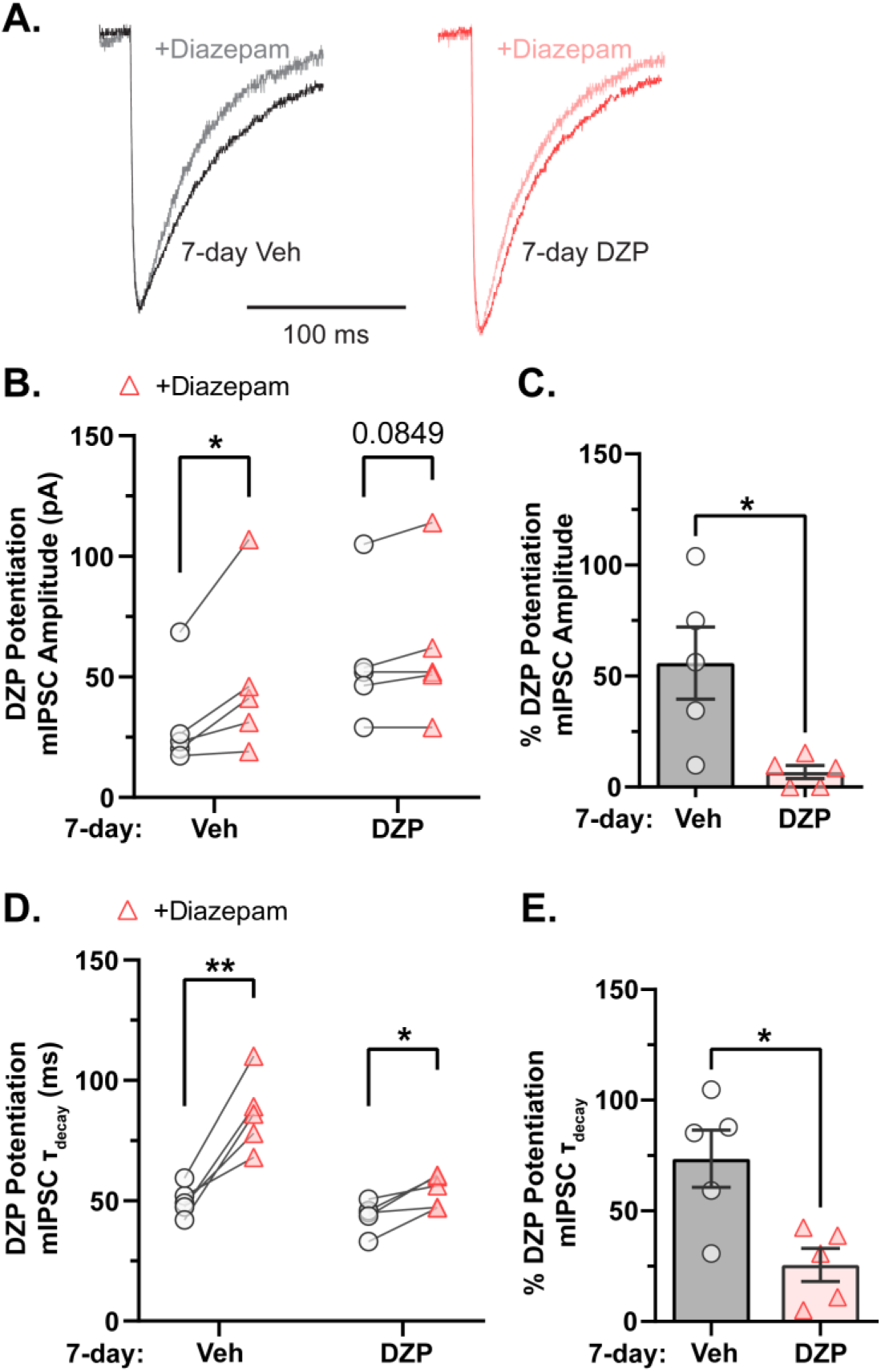
GABA_A_R potentiation by BZDs is impaired after chronic DZP treatment in primary neurons. Inhibitory postsynaptic currents were measured by whole-cell electrophysiology in neurons 7-day treated with vehicle (Veh) or 1 μM DZP and acutely exposed to 1 μM diazepam. GABA_A_R sensitivity to BZD potentiation was severely diminished after chronic DZP treatment. **(A)** Representative mIPSCs from 7-day Veh- or DZP-treated neurons before and after acute application of 1 μM diazepam. **(B)** mIPSC amplitude measured before and after application of acute diazepam (Veh-treated: initial=31.1±9.47 pA, +diazepam=48.9±15.3 pA, *p*=0.0480; DZP-treated: initial=57.2±12.7 pA, +diazepam=61.6±14.2 pA, *p*=0.0849). **(C)** The percent potentiation of mIPSC amplitude by application of acute diazepam is lost in chronic DZP-treated neurons (Veh=55.9±16.2%, DZP=6.79±2.96%; *p*=0.0176). **(D)** mIPSC τ_decay_ measured in 7-day Veh- or DZP-treated neurons before and after acute application of diazepam (Veh-treated: initial=50.0±2.87 ms, +diazepam=86.2±6.97 ms, *p*=0.0042; DZP-treated: initial=43.7±2.92 ms, +diazepam=54.3±3.00 ms, *p*=0.0197). **(E)** The percent potentiation of mIPSC τ_decay_ by acute diazepam is significantly diminished in chronic DZP-treated neurons (Veh=72.5±12.9%, DZP=25.6±7.48%; *p*=0.0125). *n*=5 cells per treatment, N=3 independent cultures; mean ± SEM. B,D: paired *t*-test; C,E: unpaired *t*-test; *\*p*≤0.05, *\*\*p*≤0.01.

### Section 3.2: Subsynaptic reorganization of gephyrin and γ2-GABAARs induced by chronic DZP treatment

Modern super-resolution microscopy has revealed synaptic proteins to be heterogeneously distributed into high-density protein clusters called subsynaptic domains (SSDs; MacGillavry et al., 2013; Nair et al., 2013; Specht et al., 2013; Crosby et al., 2019). SSDs facilitate efficient synaptic transmission and are subject to activity-dependent plasticity in response to altered neuronal activity or excitation/inhibition dysfunction (Dani et al., 2010; Specht et al., 2013; Tang et al., 2016; Pennacchietti et al., 2017; Werner et al., 2021; Yang and Annaert, 2021; Garcia et al., 2021). We hypothesized that chronic BZD treatment would disrupt the inhibitory synaptic nanoscale architecture and alter gephyrin and GABA_A_R subsynaptic organization. To this end, we employed DNA Points Accumulation for Imaging in Nanoscale Topography (DNA-PAINT), a localization-based super-resolution microscopy technique. Protein localizations are achieved by capturing and compiling many fluorescent blinking events produced by the transient binding of short single strands of DNA coupled to secondary nanobodies. Complementary DNA strands are coupled to a fluorescent dye, allowing targeting to endogenous proteins of interest and achieving protein localization with high spatial resolution (**Fig. 2A**; ^S^chnitzbauer et al., 2017^)^. Using antibodies against gephyrin and an extracellular epitope of the γ2-GABA_A_R subunit, we observed that gephyrin localizations were organized into highly concentrated clusters that aligned with vesicular GABA transporter (VGAT) puncta (identifying inhibitory presynaptic terminals) and largely overlapped with γ2-GABA_A_R localizations, while smaller clusters of both gephyrin and γ2-GABA_A_R were observed extrasynaptically (**Fig. 2B**). Assuming roughly circular SSDs and synapses, the average diameters for γ2-GABA_A_R and gephyrin synapses and SSDs (γ2-GABA_A_R, SSD: 36-40 nm, synapse: ∼210-250 nm; gephyrin, SSD: 73-82 nm, synapse: ∼360-410 nm) were within previously reported ranges (Yang and Specht, 2019; Anderson et al., 2023), confirming the validity of our technique.

**Figure 2.**
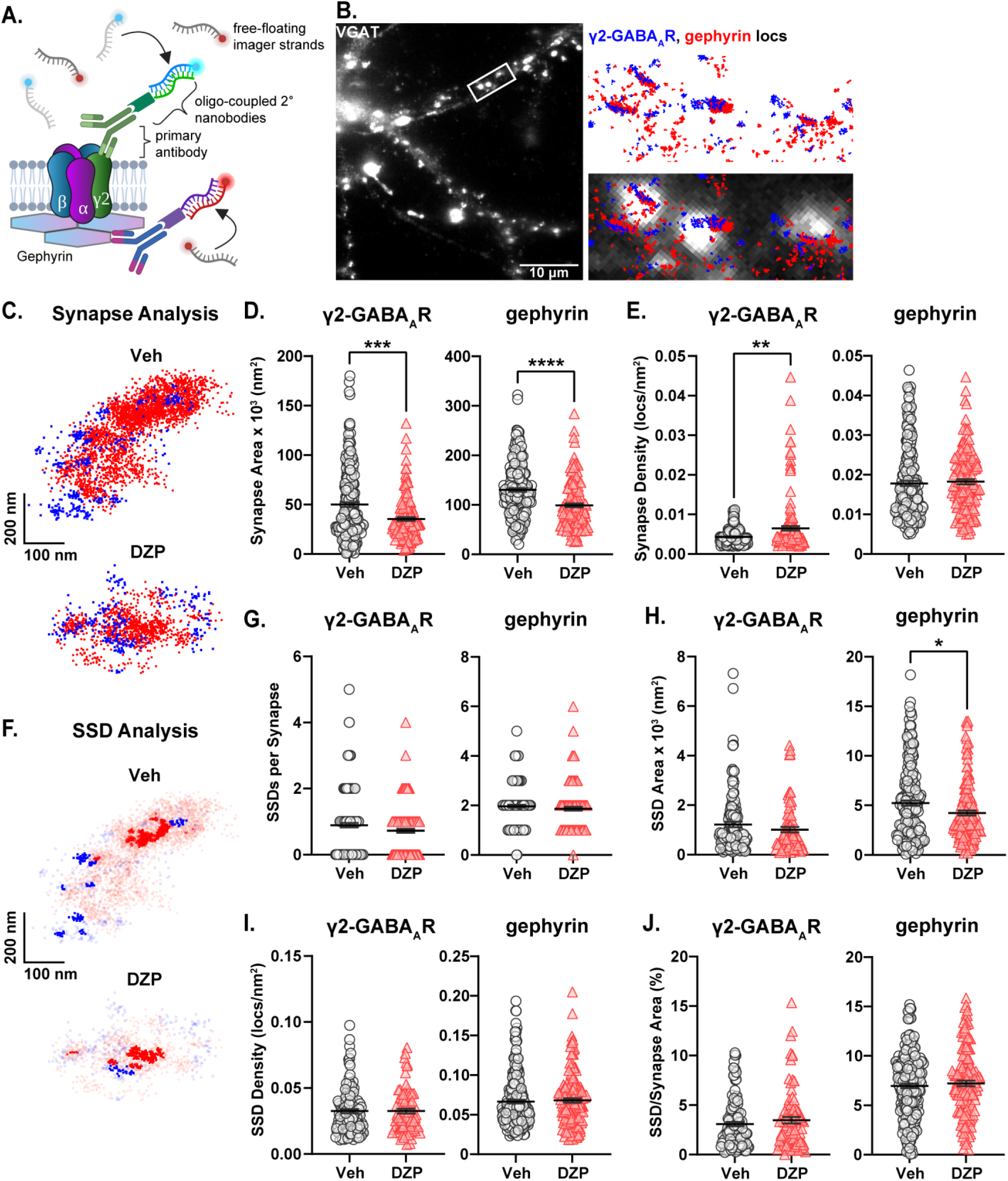
Altered subsynaptic organization of gephyrin and γ2-GABA_A_Rs by chronic DZP treatment. The subsynaptic organization of gephyrin and γ2-GABA_A_R was analyzed by DNA-PAINT super-resolution localization microscopy. **(A)** DNA-PAINT Schematic. Primary antibodies recognizing surface γ2-GABA_A_R or intracellular gephyrin are targeted by DNA-coupled secondary nanobodies (docking strands), while imager strands containing the fluorophore-bound complementary oligonucleotide remain freely available. Transient binding between the imager and docking strands produces a bright localization event. **(B)** Example snapshot of a VGAT-stained neuron used in DNA-PAINT microscopy. The zoomed region is overlayed with localizations (locs) of gephyrin (red) and γ2-GABA_A_R (blue). Colocalization with VGAT confirmed high-density localization clusters as synaptic. **(C)** Representative synapses from a 7-day Veh- or DZP-treated neuron captured by DNA-PAINT; each point represents a localization of either surface γ2-GABA_A_R (blue squares) or gephyrin (red circles). **(D,E)** Localization analysis of γ2-GABA_A_R and gephyrin total synapse area (D) and localization density (E). **(D)** Chronic DZP treatment reduced the total synapse area of both γ2-GABA_A_R (Veh=50.1±2.4×10^3^ nm^2^, DZP=35.4±1.8×10^3^ nm^2^, *p*=0.0005) and gephyrin (Veh=130.1±3.7×10^3^ nm^2^, DZP=98.9±3.9×10^3^ nm^2^; *p*<0.0001). **(E)** γ2-GABA_A_R synapse localization density was increased in DZP-treated neurons (Veh=4.3±0.13×10^-3^locs/nm^2^, DZP=6.5±0.56×10^-3^ locs/nm^2^; *p*=0.0019). Gephyrin synapse localization density was unchanged by DZP treatment. **(F)** Representative synapses from (C) with SSD localizations highlighted. **(G-J)** Analysis of γ2-GABA_A_R and gephyrin SSD numbers per synapse (G), SSD area (H), localization density within SSDs (I), and SSD/Synapse Area (J). Gephyrin SSD area was reduced after chronic DZP treatment (Veh=5.2±0.26×10^3^ nm^2^, DZP=4.2±0.25×10^3^ nm^2^; *p*=0.0265); SSDs were otherwise similar between vehicle- and DZP-treated neurons. *n*=7-9 cells, N=2 independent cultures; mean ± SEM. D,E,G-I: Mann-Whitney test, J: Mann-Whitney test (γ2-GABA_A_R) or unpaired *t*-test (gephyrin). *\*p*≤0.05, *\*\*p*≤0.01, *\*\*\*p*≤0.001, *\*\*\*\*p*≤0.0001.

Chronic DZP treatment resulted in shrinkage of the inhibitory postsynaptic area, reducing the total synapse area of γ2-GABA_A_R from ∼50 x 10^3^ nm^2^ to ∼35 x 10^3^ nm^2^ and gephyrin from ∼130 x 10^3^ nm^2^ to ∼99 x 10^3^ nm^2^ (**Fig. 2C,D**). γ2-GABA_A_Rs reorganized at higher density within this smaller area without overall loss of receptors, as indicated by a significant increase in γ2-GABA_A_R synaptic localization density (**Fig. 2E**).

Conversely, gephyrin localization density was unchanged, suggesting that chronic DZP treatment reduced total synaptic gephyrin levels (**Fig. 2E**). Consistent with this, gephyrin SSD area was also reduced by DZP treatment (**Fig. 2F,H**). However, 7-day DZP treatment did not alter the number of SSDs per synapse for either gephyrin or γ2-GABA_A_R (**Fig. 2F,G**), and γ2-GABA_A_R SSD area was also unchanged (**Fig. 2F,H**). These data indicate that chronic DZP treatment triggers a nanoscale redistribution of gephyrin and γ2-GABA_A_Rs without severely disrupting the inhibitory synaptic architecture. This is further supported by similar SSD localization density (**Fig. 2I**) and SSD/total synapse area ratios (**Fig. 2J**) between vehicle- and DZP-treated neurons for both gephyrin and γ2-GABA_A_R. Finally, to determine whether the apparent loss of synaptic gephyrin altered its alignment with GABA_A_Rs, we calculated the enrichment index for γ2-GABA_A_R and gephyrin, which is greater than one when the positioning of two proteins is closely correlated (Chen et al., 2020; Dharmasri et al., 2024). We found that chronic DZP treatment did not substantially alter the enrichment indices (**Fig. S2**), consistent with intact gephyrin-GABA_A_R synaptic associations.

### Section 3.3: Chronic DZP treatment promotes gephyrin phosphorylation and proteolytic cleavage

To investigate potential mechanisms by which chronic DZP treatment reduces synaptic gephyrin, we examined gephyrin phosphorylation at Ser270, which regulates gephyrin cluster size (Tyagarajan et al., 2011, 2013) and is increased after short-term (24 hour) DZP treatment (Lorenz-Guertin et al., 2019). Gephyrin phosphorylation was assessed using *in situ* proximity ligation (PL) assay (PLA), a novel fluorescence-based technique which detects protein-protein interactions or protein modifications with improved sensitivity and reduced nonspecific signals over traditional phospho-antibody immunofluorescence (IF). PL signals are produced when two oligonucleotide-coupled secondary antibodies (PL probes) are within close proximity (<40 nm), resulting in oligonucleotide hybridization that is then amplified and visualized by confocal microscopy as discrete, quantifiable dots (Weibrecht et al., 2010). Here, we performed PLA using an anti-gephyrin mAb7a antibody, specific for phospho-Ser270, paired with a total anti-gephyrin (3B11) antibody (**Fig. 3A**). MAP2 and VGAT counterstaining were used to identify neuronal dendrites and inhibitory synapses, respectively (**Fig. 3B**). As a control, we confirmed that minimal PL signal was observed under conditions of either primary antibody alone or with no primary antibodies (**Fig. S3**). In 7-day DZP-treated neurons, we observed trends consistent with an increase in the number of whole-field PL signals (**Fig. 3C**; *p* = 0.0757), the number of synaptic PL signals (**Fig. 3D**; *p* = 0.0751), and whole-field PL signal intensity (**Fig. 3E**; *p* = 0.0877), while synaptic PL signal intensity was significantly increased by ∼60% after chronic DZP treatment (**Fig. 3F**). These data indicate that gephyrin Ser270 phosphorylation is increased at inhibitory synapses in neurons following 7-day DZP treatment.

**Figure 3.**
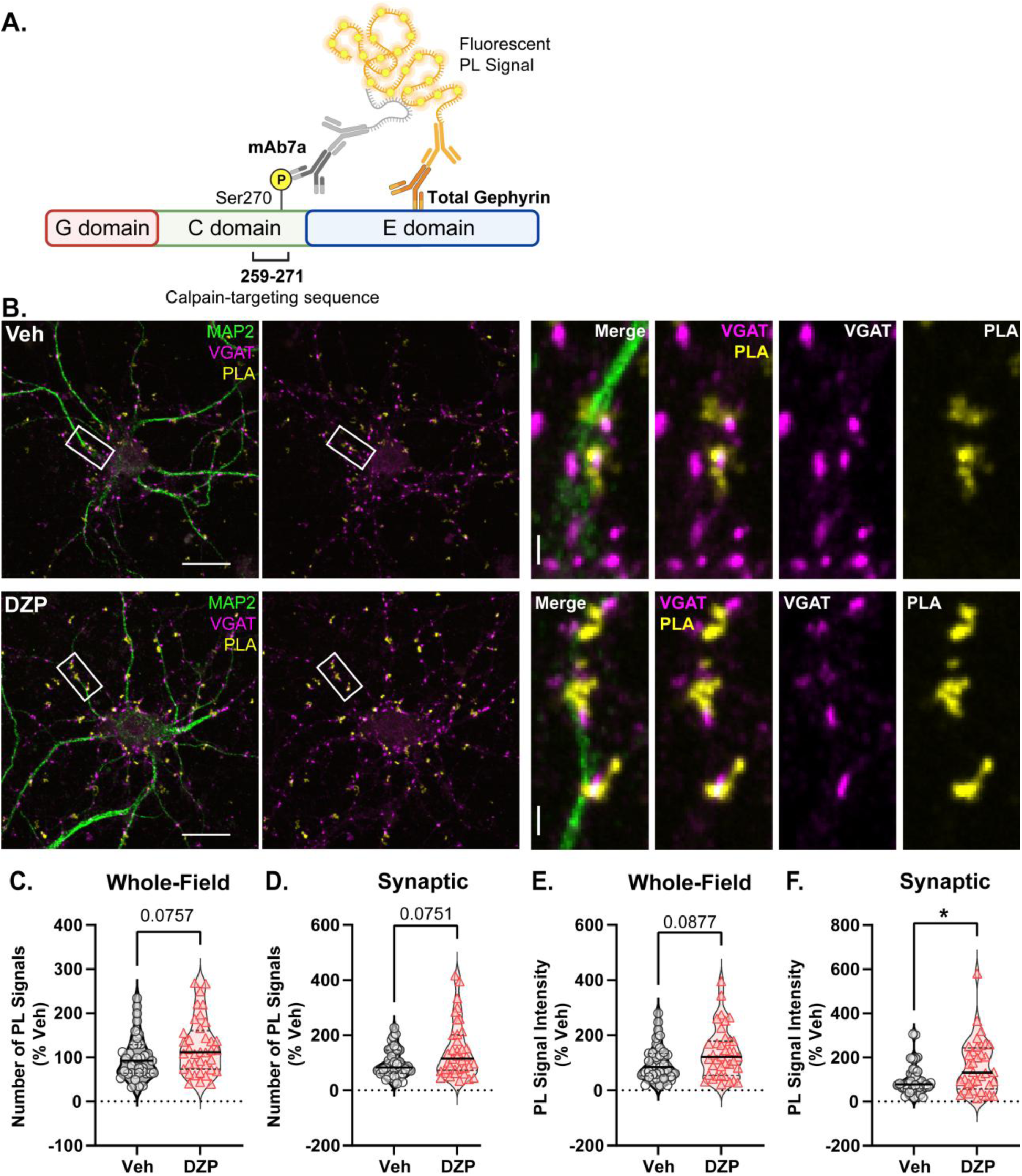
Chronic DZP treatment increases gephyrin Ser270 phosphorylation at synapses. Phosphorylation of gephyrin at Ser270 was assessed by proximity ligation (PL) assay (PLA) in 7-day Veh-versus DZP-treated neurons. **(A)** PLA Schematic; fluorescent PL signal (yellow) is only observed when the phospho-Ser270-specific mAb7a antibody is within 40 nm of the total gephyrin antibody, indicating Ser270 phosphorylation. **(B)** Representative images of Veh- or chronic DZP-treated neurons with PLA signals (yellow); MAP2 (green) and VGAT (pink) counterstaining were used to label neuronal dendrites and inhibitory synapses, respectively. **(C-F)** Quantification of the number (C,D) or intensity (E,F) of mAb7a–gephyrin PL signals in the whole field or at synaptic sites. Chronic DZP treatment significantly increased synaptic PL signal intensity, indicating increased gephyrin Ser270 phosphorylation (Veh=100.0±11.6%, DZP=157.8±20.0%; *p*=0.0263). *n*=36-37 cells, N=3 independent cultures; median (solid line) and quartiles (dashed lines) are shown. C-F: Mann-Whitney test; *\*p*≤0.05. Scale bars are 20 μm for neurons and 2 μm for dendrite zoom images.

Ser270 phosphorylation increases gephyrin susceptibility to calpain-mediated cleavage and proteolysis (Tyagarajan et al., 2011). Therefore, we next assessed chronic DZP-induced alterations to full-length and cleaved gephyrin expression using a biochemical fractionation technique followed by downstream western blotting (**Fig. 4A**). Integrity of the isolated synaptic membrane, extrasynaptic membrane, and total protein fractions was validated by immunoblotting with several synaptic and extrasynaptic markers (**Fig. S4**). Chronic DZP treatment reduced full-length gephyrin expression in the total fraction to only ∼80% that of vehicle-treated neurons (**Fig. 4B**). Consistent with our DNA-PAINT analysis (**Fig. 2**), this occurred with a near-significant decrease in synaptic full-length gephyrin (**Fig. 4B**; Veh=100.0±7.01%, DZP=75.16±9.64%; *p*=0.0559). In contrast, extrasynaptic full-length gephyrin was unchanged (**Fig. 4B**). In line with enhanced Ser270 phosphorylation, we observed three-fold higher levels of cleaved gephyrin in 7-day DZP-treated neurons (**Fig. 4C**). Surprisingly, however, this was restricted to the extrasynaptic membrane fraction (**Fig. 4C**), despite an increase in the cleaved/full-length gephyrin ratio in both the synaptic and extrasynaptic membrane fractions (**Fig. 4D**). These disparate distributions of cleaved and full-length gephyrin in the two membrane fractions may either indicate 1) a prerequisite relocation of full-length gephyrin from synaptic to extrasynaptic sites to facilitate cleavage, or 2) gephyrin cleaved at the synapse is subsequently trafficked laterally along the membrane in association with receptors, where it may accumulate in preparation for internalization and degradation.

**Figure 4.**
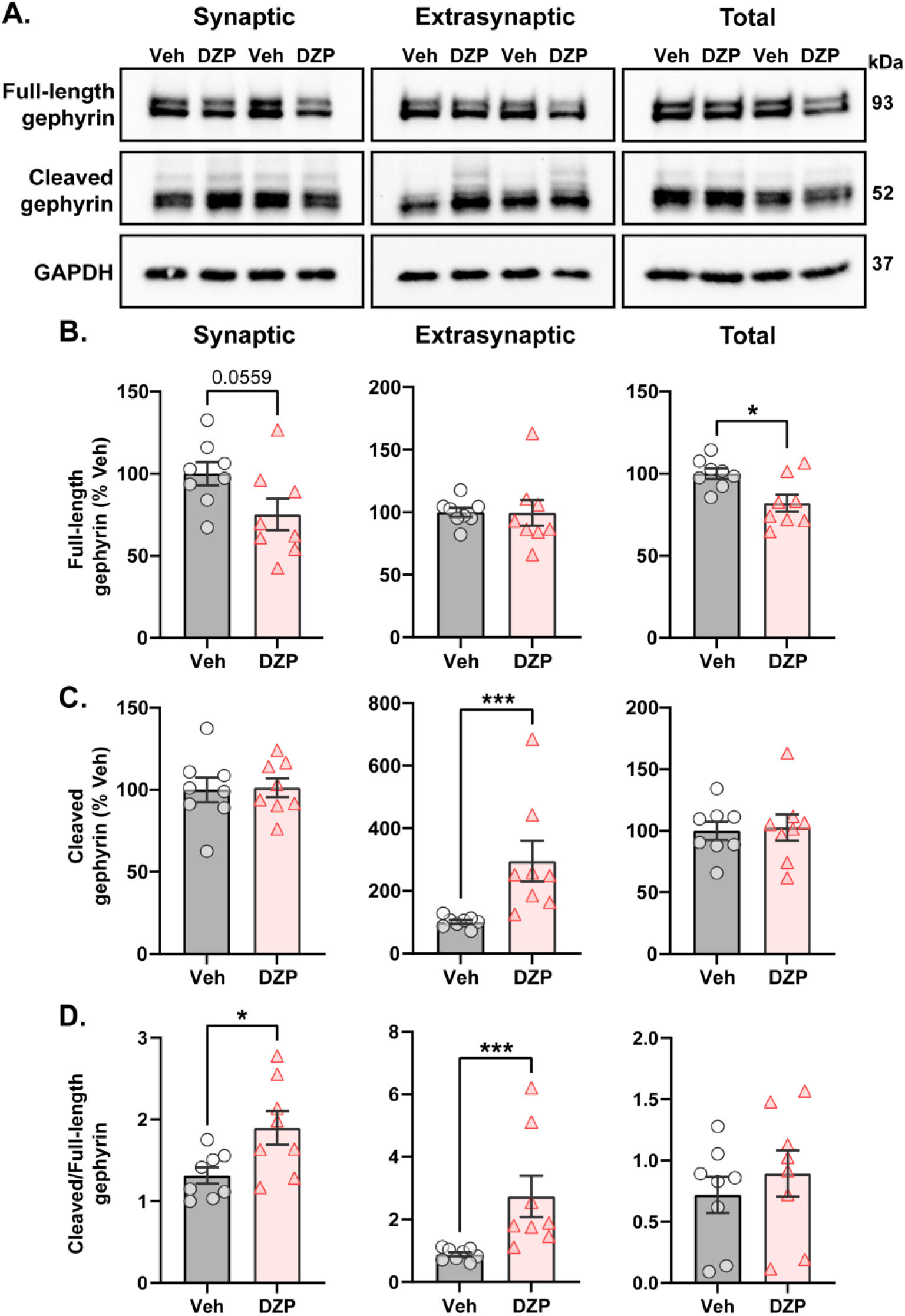
Enhanced proteolytic gephyrin cleavage decreases full-length gephyrin expression in 7-day DZP-treated neurons. Full-length or cleaved gephyrin protein expression was assessed by subcellular fractionation and western blotting. **(A)** Representative western blots. **(B-D)** Quantifications of full-length gephyrin (B), cleaved gephyrin (C), and the ratio of cleaved/full-length gephyrin (D) in the synaptic, extrasynaptic, and total protein fractions from 7-day Veh- or DZP-treated neurons. Immunoreactivities were normalized to GAPDH. **(B)** Full-length gephyrin was near-significantly reduced in the synaptic fraction (Veh=100.0±7.01%, DZP=75.16±9.64%; *p*=0.0559) and significantly reduced in the total fraction (Veh=100.0±3.14%, DZP=82.02±5.26%; *p*=0.0109), while extrasynaptic full-length gephyrin was unchanged. **(C)** Chronic DZP treatment increased cleaved gephyrin levels only at extrasynaptic sites (100.9±6.04%, DZP=295.0±65.19%, *p*=0.0003). **(D)** The proportion of cleaved/full-length gephyrin was elevated by chronic DZP treatment in the synaptic and extrasynaptic fractions (synaptic: Veh=1.31±0.099, DZP=1.90±0.20, *p*=0.0220; extrasynaptic: Veh=0.89±0.19, DZP=2.74±1.87, *p*=0.0003). *n*=2 replicates per treatment from N=4 independent cultures; mean ± SEM. B-D: unpaired *t*-test or Mann-Whitney test; *\*p*≤0.05, *\*\*\*p*≤0.001.

### Section 3.4: DZP-induced membrane redistribution of γ2-GABAARs without loss of surface or total protein expression

As loss of synaptic gephyrin can impair GABA_A_R synaptic clustering (Jacob et al., 2005; van Zundert et al., 2005; Yu et al., 2007; Carricaburu et al., 2024), we next used IF to examine γ2-GABA_A_R surface expression and subcellular localization after chronic DZP treatment. Following 7-day vehicle or DZP treatment, neurons were fixed and surface stained for γ2-GABA_A_Rs, then permeabilized and stained for the presynaptic GABA-producing enzyme, GAD65 (**Fig. 5A**). γ2-GABA_A_Rs were considered synaptic when colocalized with GAD65; otherwise, the signal was considered extrasynaptic. Chronic DZP treatment reduced the dendritic clustering density of synaptic γ2-GABA_A_Rs (**Fig. 5B**) without loss of GAD65 clusters (**Fig. 5E**), indicating a reduced proportion of inhibitory synapses expressing BZD-sensitive GABA_A_Rs. DZP treatment also decreased the γ2-GABA_A_R area per synapse without altering signal intensity (**Fig. 5B**). This is consistent with similar γ2-GABA_A_R numbers contained within a smaller postsynaptic area per synapse, in agreement with our DNA-PAINT results (**Fig. 2**). Concurrent with the loss of synaptic clusters, γ2-GABA_A_Rs were enriched extrasynaptically to 163% that of vehicle after chronic DZP treatment (**Fig. 5C**). This occurred without change to surface γ2-GABA_A_R expression (**Fig. 5D**), which was confirmed by complementary surface biotinylation analysis (**Fig. 5F,G**). These findings therefore suggest that DZP treatment induces a redistribution of synaptic γ2-GABA_A_Rs to extrasynaptic sites without altering surface expression. In contrast to short-term BZD treatments which promote γ2-GABA_A_R internalization and degradation (Nicholson et al., 2018; Lorenz-Guertin et al., 2019), biochemical analysis here additionally revealed similar total protein levels of γ2-GABA_A_R subunits in 7-day vehicle- and DZP-treated neurons (**Fig. 5F,G**).

**Figure 5.**
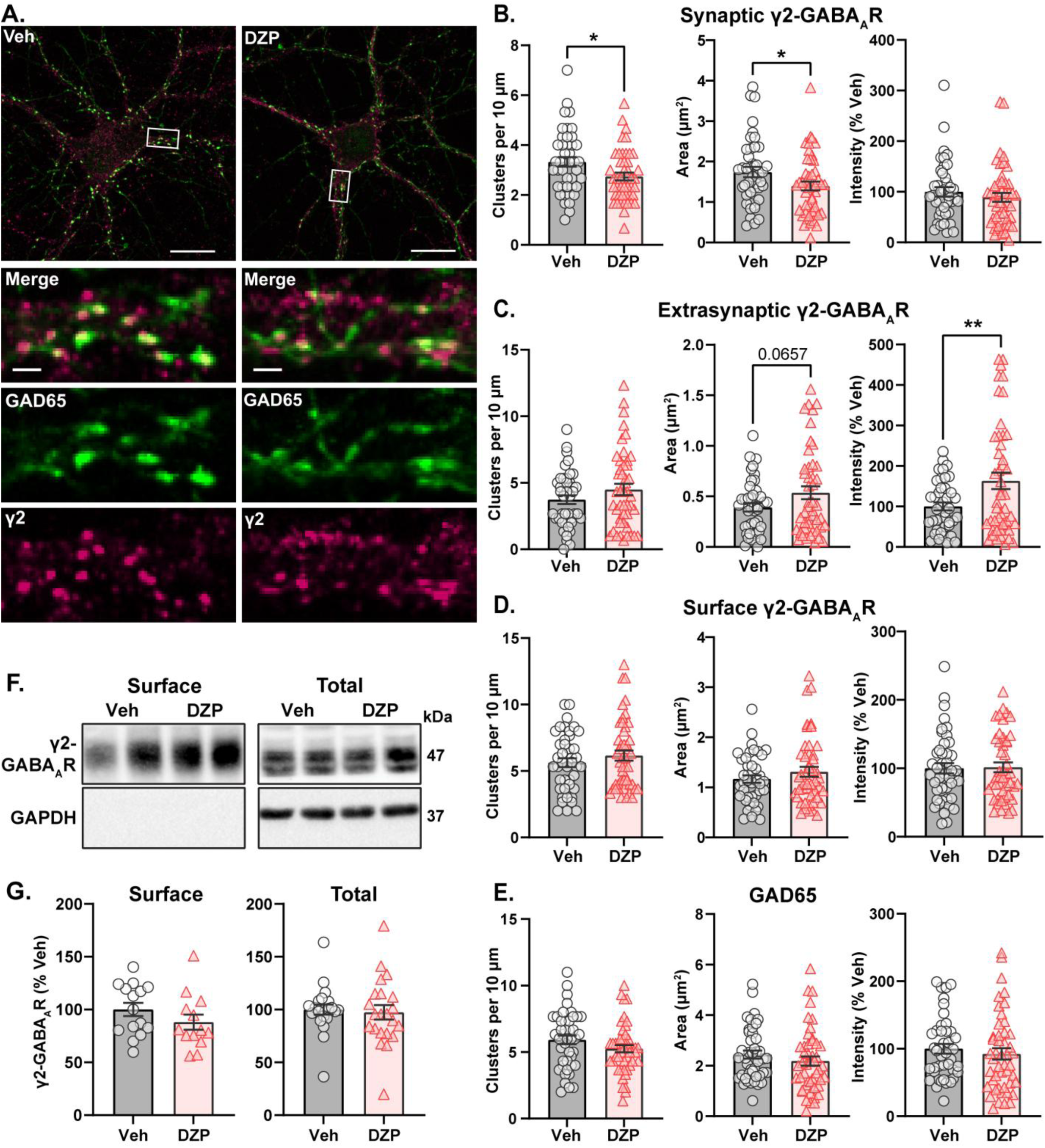
BZD-sensitive γ2-GABA_A_Rs are redistributed from synaptic to extrasynaptic sites after chronic DZP treatment without loss of surface expression. Surface expression and synaptic or extrasynaptic localization of γ2-GABA_A_Rs was assessed in 7-day Veh- or DZP-treated neurons. **(A)** Representative immunofluorescence (IF) images. Cells were first surface stained for endogenous γ2-GABA_A_R then subsequently permeabilized and stained for GAD65 to mark presynaptic inhibitory terminals. **(B-E)** Quantification of IF results, including cluster density, signal area (μm^2^), and signal intensity (% Veh) of synaptic γ2-GABA_A_Rs (B), extrasynaptic γ2-GABA_A_Rs (C), total surface γ2-GABA_A_Rs (D), or GAD65 (E). Synaptic γ2-GABA_A_R signal was defined by binary intersection with GAD65. **(B)** Chronic DZP treatment reduced γ2-GABA_A_R clustering density (Veh=3.33±0.192, DZP=2.74±0.155; *p*=0.0196) and synaptic area (Veh=1.74±0.126 μm^2^, DZP=1.40±0.109 μm^2^; *p*=0.0411) without altering signal intensity. **(C**) Chronic DZP treatment enriched the extrasynaptic accumulation of γ2-GABA_A_Rs (binary area: Veh=0.391±0.041 μm^2^, DZP=0.536±0.064 μm^2^, *p*=0.0657; sum intensity: Veh=100.0±9.66%, DZP=163.1±20.4%, *p*=0.0091). **(D-E)** Total **s**urface γ2-GABA_A_R and GAD65 staining were unchanged by DZP treatment. **(F,G)** Surface biotinylation experiments confirm that chronic DZP treatment does not alter surface or total protein expression of γ2-GABA_A_R subunits. **(F)** Representative western blots of the surface and total fractions collected by surface biotinylation. The lack of GAPDH signal in the surface fraction confirms isolation of surface proteins. **(G)** Quantification of γ2-GABA_A_R subunit surface and total protein expression. B-E: *n*=42-47 cells, N=3 independent cultures. G: *n*=15-22 replicates, N=6-9 independent cultures; mean ± SEM. B-E,G: unpaired *t*-test; *\*p*≤0.05, *\*\*p*≤0.01. Scale bars are 20 μm for neuron images and 2 μm for dendrite zoom images.

Given that mIPSC amplitude is preserved in chronic DZP-treated neurons (**Fig. S1**) despite reduced γ2-GABA_A_R cluster density, there may be a compensatory increase in the synaptic expression of non-γ2-GABA_A_Rs. BZD-insensitive α4-GABA_A_Rs are elevated in some neurodevelopmental disorders and are associated with BZD-resistant seizures (Talos et al., 2012; Sharma et al., 2021). Therefore, we assessed α4-GABA_A_R synaptic and total subunit expression in neurons after chronic DZP treatment. While α4-GABA_A_R synaptic levels were similar between vehicle- and DZP-treated neurons, total protein expression trended upward (**Fig. S5**; Veh=100±10%, DZP=144±18%, *p* = 0.0623). Together, these data are consistent with a model of individual synapse-specific losses of BZD-sensitive γ2-GABA_A_Rs induced by chronic DZP treatment and gephyrin downregulation which are not sufficiently widespread to reduce mIPSCs and/or are compensated for by upregulation of other receptor subtypes.

### Section 3.5: Gephyrin and γ2-GABAAR interactions and trafficking dynamics are altered by chronic DZP treatment

To determine whether GABA_A_R accumulation in the extrasynaptic membrane and reduced synaptic clustering is mediated by impaired gephyrin-GABA_A_R interactions, we again employed PLA and paired a γ2-GABA_A_R antibody with a total gephyrin (3B11) antibody (**Fig. 6A**). PLA was performed under permeabilized conditions and thus included gephyrin-GABA_A_R interactions both at the cell surface and intracellularly. As before, MAP2 and VGAT counterstaining was used to identify neuronal dendrites and inhibitory synapses, respectively (**Fig. 6B**). Unexpectedly, chronic DZP treatment produced a near-significant increase in the total number of whole-field gephyrin-GABA_A_R PL signals (**Fig. 6C**; *p* = 0.0561), consistent with enhanced receptor-scaffold associations. Stratifying the PL signals into synaptic or extrasynaptic by colocalization with VGAT revealed similar numbers of synaptic PL signals between vehicle- and DZP-treated neurons (**Fig. 6D**). Surprisingly, however, chronic DZP treatment produced higher numbers of extrasynaptic gephyrin-GABA_A_R PL signals (**Fig. 6E**). Therefore, these findings suggest that interactions of gephyrin with BZD-sensitive GABA_A_Rs are elevated specifically at extrasynaptic sites following chronic DZP treatment.

**Figure 6.**
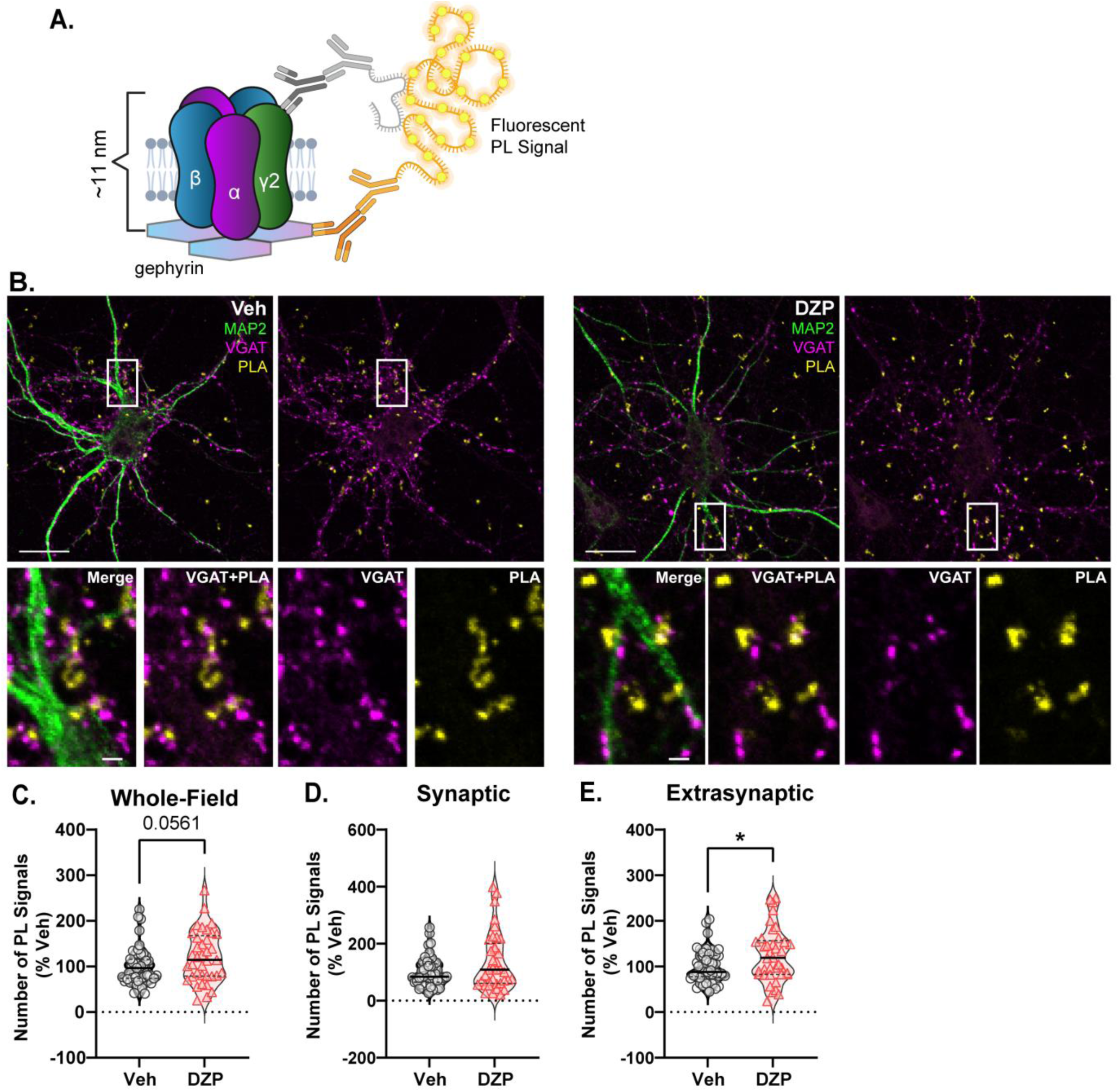
Gephyrin associations with γ2-GABA_A_Rs are enhanced by chronic DZP treatment in the extrasynaptic membrane. Gephyrin associations with γ2-GABA_A_Rs were assessed by PLA in neurons treated with Veh or 1 μM DZP for 7 days. **(A)** PLA Schematic; fluorescent PL signal is only observed when γ2-GABA_A_R and gephyrin are within 40 nm, indicating association; experiments were performed under permeabilized conditions. **(B)** Representative neuron images; MAP2 and VGAT counterstaining was included to label neuronal dendrites and inhibitory synapses, respectively; PLA signals are shown in yellow. **(C-E)** Quantification of the number of whole-field (C), synaptic (D), or extrasynaptic (E) gephyrin-GABA_A_R PL signals. Chronic DZP treatment resulted in a near-significant increase in the number of whole-field PL signals (C; Veh=100.0±6.015%, DZP=121.1±9.219%; *p*=0.0561) and significantly higher numbers of extrasynaptic PL signals (E; Veh=100.0±5.328%, DZP=125.0±9.508%; *p*=0.0270), while synaptic PL signals were similar between vehicle- and DZP-treated neurons. *n*=38-47 cells, N=3 independent cultures; median (solid line) and quartiles (dashed lines) are shown. C-E: Mann-Whitney test; *\*p*≤0.05, *\*\*p*≤0.01. Scale bars are 20 μm for neurons and 2 μm for dendrite zoom images.

The preservation of synaptic gephyrin-GABA_A_R associations (**Figs. 6,S2**) may suggest that synaptic stability is intact even during chronic DZP treatment, which is conversely impaired after 12-24 hour DZP exposure (Vlachos et al., 2013; Lorenz-Guertin et al., 2019). On the other hand, extrasynaptic gephyrin interactions with glycine receptors can slow their membrane diffusion (Ehrensperger et al., 2007). Thus, we hypothesized that the increase in extrasynaptic gephyrin-GABA_A_R interactions (**Fig. 6**) may similarly slow extrasynaptic γ2-GABA_A_Rs, potentially facilitating their extrasynaptic accumulation (**Fig. 5**). To assess trafficking dynamics, we performed live-cell FRAP (fluorescence recovery after photobleaching) experiments in hippocampal neurons co-transfected with mScarlet-gephyrin.FingR and γ2^pH^FAP constructs. mScarlet-gephyrin.FingR is a transcriptionally controlled fibronectin intrabody generated with mRNA display (FingR) that selectively binds endogenous gephyrin without impacting protein levels or synaptic architecture (Gross et al., 2013; Bensussen et al., 2020). The γ2^pH^FAP subunit construct has an extracellular pH-sensitive pHluorin tag, allowing surface-specific fluorescence, and a fluorogen-activating peptide (FAP) tag that binds malachite green (MG) dyes with high specificity. γ2^pH^FAP assembles with endogenous subunits into receptors that show normal GABA response, DZP potentiation, and trafficking (Lorenz-Guertin et al., 2017, 2019).

We first confirmed that the majority of mScarlet-gephyrin.FingR clusters were synaptic by live labeling of inhibitory presynaptic terminals with a fluorescently tagged antibody to VGAT (VGAT CypHer5E; **Fig. S6A**), which was added to the neuron dish for 1-2 hours to allow uptake into synaptic vesicles. In agreement with previously reported values of ∼DIV 21 neurons (Danglot et al., 2003), approximately 90% of analyzed gephyrin clusters were colocalized with VGAT (**Fig. S6B**). Synaptic γ2^pH^FAP signals were thus subsequently defined by colocalization with bright clusters of mScarlet-gephyrin.FingR. Following an initial pre-bleach acquisition phase to establish baseline fluorescence, we photobleached synaptic (**Fig. 7A**) and extrasynaptic (**Fig. 7C**) regions of neurons expressing γ2^pH^FAP and mScarlet-gephyrin.FingR. Fluorescence recovery within these regions was monitored every two minutes for the next 30 minutes. MG-βTau, a cell-impermeable MG dye that is nonfluorescent until FAP binding (Yan et al., 2015), was added immediately after photobleaching to confirm surface expression of the γ2^pH^FAP GABA_A_R clusters (**Fig. S6C,D**). Chronic DZP treatment resulted in higher fluorescence recovery of synaptic mScarlet-gephyrin.FingR (**Fig. 7A,B**) compared to vehicle-treated neurons. This is consistent with enhanced synaptic gephyrin turnover and a reduction in the population of stable, immobilized gephyrin at synapses. Despite this destabilization of gephyrin, the synaptic dynamics of γ2-GABA_A_R were unchanged by chronic DZP treatment (**Fig. 7B**). In agreement with our hypothesis, the fluorescence recovery of γ2^pH^FAP was significantly lower in extrasynaptic regions of DZP-treated neurons (**Fig. 7C,D**), while extrasynaptic mScarlet-gephyrin.FingR trafficking was unchanged (**Fig. 7D**). Because GABA_A_Rs are primarily exocytosed extrasynaptically (Bogdanov et al., 2006), yet surface γ2-GABA_A_R expression was unchanged by DZP treatment (**Fig. 5**), these data are consistent with reduced lateral receptor movements within the extrasynaptic membrane rather than decreased forward trafficking or altered diffusion from synaptic to extrasynaptic sites. These findings collectively suggest that distinct alterations in the trafficking dynamics of gephyrin and γ2-GABA_A_Rs may effectively diminish BZD sensitivity by reducing the synaptic prevalence of BZD-sensitive receptors and restricting their movements extrasynaptically.

**Figure 7.**
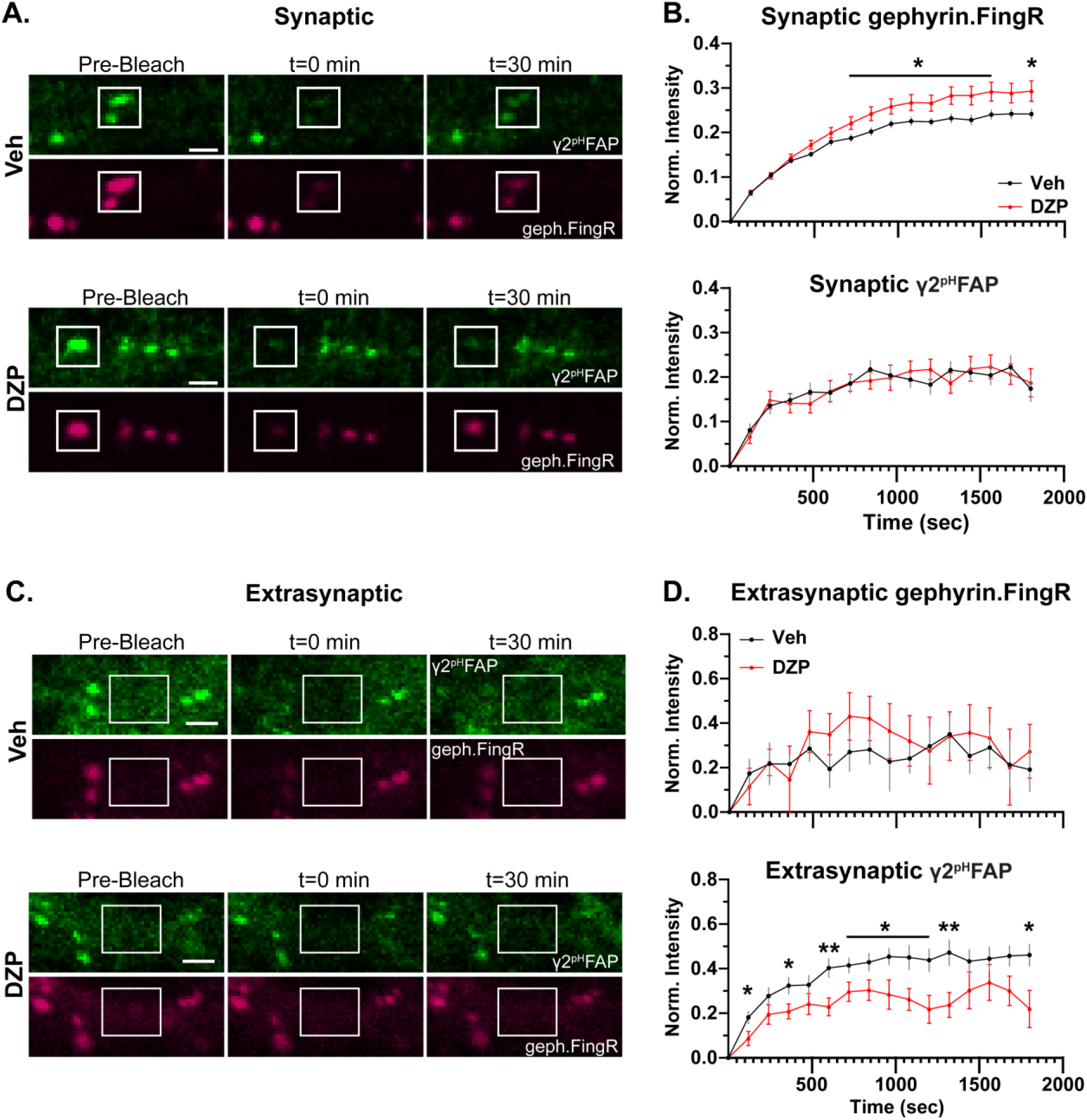
Chronic DZP treatment destabilizes synaptic gephyrin and impairs the mobility of extrasynaptic γ2-GABA_A_Rs. Fluorescence Recovery After Photobleaching (FRAP) experiments were performed in hippocampal neurons co-transfected with γ2^pH^FAP and mScarlet-Gephyrin.FingR and treated with Veh or 1 μM DZP for 7 days. **(A,C)** Representative images of synaptic (A) or extrasynaptic (C) γ2^pH^FAP and mScarlet-Gephyrin.FingR (geph.FingR, gephyrin.FingR) before bleaching (pre-bleach), immediately after bleaching (t=0 min), and 30 min post-bleach (t=30 min). White boxes indicate regions of photobleaching. **(B,D)** Fluorescence recovery was monitored after photobleaching every two minutes for 30 minutes for both synaptic (B) or extrasynaptic (D) regions. **(B)** mScarlet-gephyrin.FingR synaptic turnover was elevated by chronic DZP treatment, while synaptic γ2^pH^FAP recovery was unchanged by DZP treatment. **(D)** Chronic DZP treatment did not affect extrasynaptic trafficking of mScarlet-gephyrin.FingR, while γ2^pH^FAP extrasynaptic mobility was reduced. B,D: *n*=16-17 cells, N=3 independent cultures; mean ± SEM. Analyses by multiple unpaired *t*-tests; *\*p*≤0.05, *\*\*p*≤0.01. Scale bars are 20 μm for neurons and 2 μm for dendrite zoom images.

## Section 4: Discussion

BZD clinical use is severely limited by the rapid development of tolerance to the therapeutic effects. This can drive a need for dose escalation that increases risk of patient abuse, addiction, and dependence associated with a withdrawal syndrome that occurs upon drug discontinuation marked by sleep disturbance, anxiety, panic attacks, and other neurological hallmarks of impaired inhibition (Pétursson, 1994; Janhsen et al., 2015). While many studies have described initial neuronal adaptations after acute or short-term BZD exposure, there has been a lack of research focused on long-term neuroplasticity mechanisms underlying BZD tolerance. With as many as 25% of all BZD users continuing use for several months to years at a time (Olfson et al., 2015; Kurko et al., 2015; Kaufmann et al., 2018; Tanguay Bernard et al., 2018) and the 50% rates of relapse following BZD discontinuation (Morin et al., 2005; Gerlach et al., 2019; Chapoutot et al., 2021), there is an urgent need to understand the impact of long-term BZD treatments on GABA_A_R regulation and inhibitory synapse plasticity.

In this study, we describe key alterations to the inhibitory postsynaptic scaffold gephyrin and BZD-sensitive γ2-GABA_A_Rs in primary neurons chronically treated with DZP. Following functional confirmation of diminished BZD sensitivity (**Fig. 1**), we provide the first analysis of BZD-induced changes to inhibitory subsynaptic organization using super-resolution DNA-PAINT localization microscopy. For gephyrin, DNA-PAINT analysis found a DZP-induced decrease in total and subsynaptic domain area (**Fig. 2**). A loss of synaptic and total gephyrin protein expression was then confirmed by biochemical fractionation analysis (**Fig. 4**). Furthermore, this was associated with increased gephyrin Ser270 phosphorylation (**Fig. 3**), protease-mediated gephyrin cleavage (**Fig. 4**), and impaired synaptic gephyrin stability (**Fig. 7**). These results collectively demonstrate that chronic DZP treatment activates signaling pathways which promote the deconstruction of this critical inhibitory scaffold. As gephyrin regulates GABA_A_R clustering, we also assessed DZP-induced changes to GABA_A_Rs.

Super-resolution analysis revealed increased γ2-GABA_A_R localization density per synapse (**Fig. 2**). Corroborated by immunofluorescence (**Fig. 5**), this is consistent with γ2-GABA_A_Rs clustering within a smaller postsynaptic area without loss of receptors per synapse. However, there were overall fewer inhibitory synapses expressing γ2-GABA_A_Rs (**Fig. 5**). Since presynaptic GAD65 (**Fig. 5**) and basal mIPSC parameters (**Fig. S1**) were preserved, this indicates that chronic DZP treatment reduced the proportion of inhibitory synapses containing BZD-sensitive GABA_A_Rs. Rather than being removed from the cell surface, γ2-GABA_A_Rs were enriched extrasynaptically after chronic DZP treatment (**Fig. 5**). Interestingly, the lateral mobility of these extrasynaptic receptors was restricted in DZP-treated neurons (**Fig. 7**), which we demonstrate by PLA to correlate with higher levels of gephyrin-GABA_A_R associations away from the synapse (**Fig. 6**). In summary, these findings uncover important plasticity mechanisms of gephyrin and γ2-GABA_A_Rs during extended BZD treatment. We propose that these processes together limit the synaptic prevalence and renewal of BZD-sensitive GABA_A_Rs to chronically diminish synaptic sensitivity to BZDs without substantially impairing inhibitory neurotransmission.

Postsynaptic receptors and scaffolds at synapses form small (<100 nm diameter), high-density subsynaptic clusters that trans-synaptically align with active zone machinery in the presynaptic terminal, facilitating efficient neurotransmission (Tang et al., 2016; Crosby et al., 2019; Gookin et al., 2022; Olah et al., 2023). Recent work with super-resolution microscopy, particularly localization-based, has established the importance of SSDs in postsynaptic organization and plasticity (Chen et al., 2018; reviewed in Yang and Specht, 2019). However, the impact of chronic BZD treatment on the inhibitory subsynaptic organization has not been previously described. Here, DNA-PAINT studies revealed an overall shrinkage of the gephyrin and γ2-GABA_A_R synapse areas, a redistribution of synaptic γ2-GABA_A_Rs within this smaller area, and smaller gephyrin SSDs (**Fig. 2**). However, the relative timing and potential interdependence of the respective gephyrin and γ2-GABA_A_R nanoscale rearrangements remain undetermined. While some studies have suggested a largely cooperative relationship between GABA_A_Rs and gephyrin (Essrich et al., 1998; Schweizer, 2003; Alldred et al., 2005; Crosby et al., ^2^019^)^, others have described entirely independent mechanisms of receptor and scaffold plasticity (Niwa et al., 2012; Garcia et al., 2021; Merlaud et al., 2022). A highly coordinated subsynaptic relationship was demonstrated by expression of a dominant-negative gephyrin construct that disrupted both γ2-GABA_A_R and gephyrin SSD size and positioning (Crosby et al., 2019). Use of antisense oligonucleotides to block gephyrin expression has also been shown to promote a switch in the synaptic GABA_A_R population to receptors that were highly sensitive to zinc (non-γ2-GABA_A_R) and insensitive to BZDs (van Zundert et al., 2005), further suggesting a specific role for gephyrin in the insertion and stabilization of BZD-sensitive γ2-GABA_A_R clusters. On the other hand, GABA_A_R lateral diffusion from synapses in response to acute increases in neuronal activity temporally preceded that of gephyrin (Niwa et al., 2012). Similarly, during acute excitotoxic insult, calcineurin dephosphorylation of the γ2 subunit first reduced γ2-GABA_A_R SSDs, which was then followed by gephyrin cleavage and SSD disassembly (Garcia et al., 2021). Taking our FRAP data into account, the shrinkage in gephyrin SSDs is accompanied by reduced synaptic stability in DZP-treated neurons (**Figs. 2,7**). Conversely, γ2-GABA_A_R SSDs are unchanged by DZP treatment, and synaptic stability is also maintained (**Figs. 2,7**). These distinct alterations in SSD and synaptic turnover for the scaffold and receptor may indicate that synaptic stabilization of γ2-GABA_A_Rs is independent of gephyrin stability under conditions of chronic DZP treatment.

This would likely be due to additional interactions of γ2-GABA_A_R with other postsynaptic proteins, including neuroligin-2 and GARLH4 (Davenport et al., 2017; Yamasaki et al., 2017; Martenson et al., 2017). To fully understand the gephyrin-GABA_A_R relationship, future studies should perform time-course analysis of SSDs throughout the chronic BZD treatment. Additionally, experiments utilizing inhibitors to the protease calpain could determine whether gephyrin disassembly is required for γ2-GABA_A_Rs subsynaptic redistribution.

Chronic BZD treatment has been shown to produce distinct molecular responses dependent upon the brain region (Impagnatiello et al., 1996; Longone et al., 1996; Pesold et al., 1997; Wu et al., 1994; Li et al., 2000; Wright et al., 2014; Furukawa et al., 2017), method (Fernandes and File, 1999; Arnot et al., 2001; Allison and Pratt, 2006) and length (Wu et al., 1994; Holt et al., 1996; Ferreri et al., 2015) of dosing, and behavioral effect analyzed (Fernandes and File, 1999; Bateson, 2002; Vinkers and Olivier, 2012). Consequently, comparisons of prior studies of BZD tolerance are challenging due to discrepancies in treatment paradigm, BZD ligand used, and brain regions assessed. Though lacking the complexity and connectivity of an *in vivo* system, primary neuronal culture is a simplified model that readily permits high-resolution analysis of precise molecular mechanisms of plasticity, including changes to synaptic protein trafficking dynamics, intermolecular interactions, and subsynaptic organization. We previously used this system to describe the initial neuroplasticity mechanisms triggered by short-term (24 hour) DZP exposure (Lorenz-Guertin et al., 2019).

Here, we used the same primary neuronal culture system, BZD ligand, and BZD concentration while extending the length of drug treatment to discern the differential neuroplasticity induced by sustained, chronic BZD exposure. For gephyrin, we found that chronic DZP treatment resulted in a disruption of the gephyrin scaffold via altered posttranslational processing heavily reminiscent of the 24 hour phenotype (Lorenz-Guertin et al., 2019). Thus, DZP treatment produces a moderate yet persistent downregulation of synaptic gephyrin expression and stability. This long-lasting destabilization may be expected to disrupt gephyrin’s critical role in the clustering of GABA_A_Rs, but gephyrin-GABA_A_R interactions have not before been analyzed during chronic exposure to DZP. This is particularly important given that the GABA_A_R gephyrin binding domain appears conformationally linked to that of the BZD binding domain (Gouzer et al., 2014; Lévi et al., 2015). In this study, we provide the first analysis of the gephyrin-GABA_A_R association after chronic DZP treatment using PLA and surprisingly found that gephyrin interactions with γ2-GABA_A_Rs at the synapse were not reduced (**Fig. 6**). It is possible that the PLA analysis lacks sufficient resolution to discern subtle changes in association, particularly within the postsynaptic density which contains high concentrations of these proteins. Additionally, this assay does not provide more detailed information as to whether the strength or nature of the interaction is altered; thus, detailed characterization of this interaction and the consequential impact on BZD binding may be an important avenue of future research.

In contrast to gephyrin, γ2-GABA_A_Rs exhibit several neuroplasticity alterations after chronic DZP treatment that are distinct from short-term exposure. We previously showed that 24 hour DZP treatment impairs synaptic γ2-GABA_A_R stability and reduces γ2-GABA_A_R subunit expression via increased lysosomal-mediated degradation ^(^Lorenz-Guertin et al., 2019^)^. Similarly, other groups have also reported reduced expression of BZD-sensitive GABA_A_Rs (Jacob et al., 2012; Nicholson et al., 2018; Foitzick et al., 2020; González Gómez et al., 2023) and reduced mIPSCs (Jacob et al., 2012; Nicholson et al., 2018) after short-term (<72 hour) BZD treatment.

Conversely, our findings reveal that these initial adaptations in γ2-GABA_A_Rs do not persist with chronic DZP treatment, as γ2-GABA_A_R total protein and surface expression were maintained (**Fig. 5**) and mIPSCs were preserved (**Fig. S1**). Instead, γ2-GABA_A_Rs were redistributed throughout the surface membrane: there were fewer inhibitory postsynaptic sites which expressed γ2-GABA_A_Rs (**Fig. 5**), and for those that did, these receptors were condensed within a smaller area (**Figs. 2,5**), which is potentially due to the reduced overall postsynaptic and subsynaptic gephyrin area (**Fig. 2**). The trafficking dynamics of γ2-GABA_A_Rs were also distinctly impacted by chronic versus short-term DZP treatment. While 24-hour DZP treatment accelerated γ2-GABA_A_R synaptic turnover without impacting extrasynaptic dynamics (Lorenz-Guertin et al., 2019), we instead found that 7-day DZP treatment did not affect the synaptic exchange of γ2-GABA_A_Rs despite the destabilization in gephyrin (**Fig. 7**). These alterations collectively produced an accumulation of γ2-GABA_A_Rs extrasynaptically (**Fig. 5**). γ2-GABA_A_Rs represent the major synaptic GABA_A_R population (Olsen and Sieghart, ^2^008, 2009^)^, and the γ2 subunit is required for proper postsynaptic clustering maintenance (Essrich et al., 1998; Martenson et al., 2017) and organism viability (Schweizer, 2003). Hence, downregulation of this critical receptor subtype is evidently minimized in the long-term.

While synaptic γ2-GABA_A_R trafficking was maintained, extrasynaptic lateral mobility was reduced in chronic DZP-treated neurons (**Fig. 7**), which correlated with enhanced gephyrin-GABA_A_R interactions extrasynaptically (**Fig. 6**). This suggests that an extrasynaptic pool of gephyrin is restricting the diffusion of γ2-GABA_A_Rs extrasynaptically, which has been demonstrated for glycine receptors (Ehrensperger et al., 2007). This may effectively reduce the speed with which BZD-sensitive GABA_A_Rs are re-incorporated into the synapse.

However, it is also possible that these gephyrin-GABA_A_R extrasynaptic interactions are mediated by a temporary continued association of receptors with cleaved gephyrin fragments. The C-terminal gephyrin cleavage fragment, potentially including an intact receptor binding site, is relatively long-lived (Kawasaki et al., 1997). Under physiological conditions, calcium-dependent calpain proteolysis regulates gephyrin clustering and contributes to neurite outgrowth and synapse remodeling (Kawasaki et al., 1997). In contrast, calcium overload leads to excessive, pathological calpain activity (Bevers and Neumar, 2008; Vosler et al., 2008), promoting gephyrin degradation, disassembly, and a loss of synaptic γ2-GABA_A_Rs (Costa et al., 2016). The subcellular localization of cleaved gephyrin fragments has not been before described, and the order in which gephyrin is cleaved and removed from the synapse is unclear. Increased gephyrin cleavage has been observed within ∼9 minutes of oxygen-glucose deprivation (OGD) in neuron culture, but gephyrin SSD volume was not reduced until ∼15 minutes (Garcia et al., 2021). Together with our data showing enrichment of cleaved gephyrin fragments specifically in the extrasynaptic membrane, this suggests that gephyrin is first cleaved at the synapse and subsequently diffuses to extrasynaptic sites. This is likely in preparation for receptor and scaffold internalization and degradation. Interestingly, when GABA_A_Rs are in an active or desensitized conformational state, they are removed with gephyrin from the synapse, where they then localize together in extrasynaptic endocytic zones (Merlaud et al., 2022). As the PLA (**Fig. 6**) was performed under permeabilized conditions, both surface and internal gephyrin-GABA_A_R associations are included. Thus, activated γ2-GABA_A_Rs may diffuse from the synapse together with cleaved gephyrin fragments during chronic BZD treatment for internalization extrasynaptically.

Gephyrin susceptibility to calpain-mediated cleavage and proteolysis is enhanced when phosphorylated by the kinase GSK3β at Ser270, resulting in reduced gephyrin clustering (Tyagarajan et al., 2011). In accordance with the elevated cleavage levels **(Fig. 4**), we found by PLA that chronic DZP treatment increased gephyrin Ser270 phosphorylation (**Fig. 3**). Gephyrin forms a planar submembrane hexagonal lattice at synapses through trimerization and dimerization of its N-terminal G- and C-terminal E-domains, respectively. The largely disordered central linker C-domain is the main target for posttranslational modifications (PTMs), which provide control of scaffold size, stability, and packing density (Zacchi et al., 2014; Choii and Ko, 2015; Kasaragod and Schindelin, 2018; Groeneweg et al., 2018) by modifying the degree to which the C-domain is folded or extended, consequently altering the compaction of the entire scaffold lattice (Sander et al., 2013; Groeneweg et al., 2018). Alanine mutation of the Ser270 residue to block phosphorylation was revealed by localization microscopy to reduce gephyrin packing density (Battaglia et al., 2018). Thus, the reduction in gephyrin synaptic and SSD areas triggered by chronic DZP treatment (**Fig. 2**) may not only be mediated by the loss in protein expression (**Fig. 4**) but also by enhanced Ser270 phosphorylation (**Fig. 3**) and scaffold packing density.

However, evidence of crosstalk between gephyrin PTMs complicates the current understanding of scaffold regulation (Tyagarajan et al., 2013). For example, Ser270 cooperates with the ERK1/2 Ser268 site to dynamically control gephyrin clustering and proteolysis (Tyagarajan et al., 2013). Other kinases regulating gephyrin include PKA and CaMKII, which modulate gephyrin plasticity responses (Flores et al., 2015). Gephyrin is also modified by acetylation (Tyagarajan et al., 2013; Ghosh et al., 2016); S-nitrosylation (Dejanovic and Schwarz, 2014; Yang et al., 2024); palmitoylation (Dejanovic et al., 2014; Shen et al., 2019); and SUMOylation (Ghosh et al., 2016). The majority of gephyrin PTMs have not been thoroughly characterized; given the extent to which gephyrin is posttranslationally modified and the complexity of these interactions, comprehensive proteomics and PTM site mutation studies are needed to fully understand their role in BZD tolerance.

BZDs have remained important clinical drugs for decades due to their ability to mediate anxiolytic, anticonvulsant, and sedative effects with high efficacy and low toxicity. However, they are limited by the rapid development of tolerance and dependence, the mechanisms of which have remained unresolved. Here, we describe key features of inhibitory synaptic plasticity occurring in primary neurons chronically treated with DZP, including: 1) reduced synaptic expression and altered subsynaptic organization of gephyrin and γ2-GABA_A_Rs; 2) increased gephyrin Ser270 phosphorylation, proteolysis, and synaptic turnover; 3) extrasynaptic accumulation and reduced mobility of γ2-GABA_A_Rs; and 4) increased extrasynaptic associations between γ2-GABA_A_Rs and gephyrin. Collectively, these disruptions may both impair the conformational relationship between the gephyrin and BZD receptor binding sites and constrict the ability of BZD-sensitive γ2-GABA_A_Rs to return to synaptic sites following extrasynaptic dispersal. At least on a 7-day treatment timeline *in vitro*, these changes occurred without loss of baseline mIPSC parameters, presynaptic GAD65 expression, or surface and total γ2-GABA_A_R subunit protein levels. Similarly, chronic DZP treatment *in vivo* at 10 mg/kg daily dosing did not impact baseline synaptic inhibition (Lorenz-Guertin et al., 2023). However, longer treatments may result in further impairment of inhibition, and future studies *in vivo* at and beyond the 2-4 week FDA treatment guidelines are needed. Important changes to inhibitory synapses described here with long-term DZP treatment are often distinct from those observed with short-term BZD treatment (Jacob et al., 2012; Nicholson et al., 2018; Lorenz-Guertin et al., 2019), further underpinning a need for more detailed mechanistic insight during longer BZD treatments. This is especially true given the high prevalence of prolonged BZD use in patient populations (Kurko et al., 2015; Olfson et al., 2015; Kaufmann et al., 2018; Tanguay Bernard et al., 2018). Future *in vitro* and *in vivo* studies are also needed to define upstream mechanisms responsible for the described changes in γ2-GABA_A_R and gephyrin regulation with 7-day and longer DZP treatments, including: 1) examining excitatory glutamatergic receptors as sources of calcium influx and crosstalk signaling; 2) gephyrin and γ2-GABA_A_R localization to endocytic zones and internalization processes; and 3) proteomic analysis to comprehensively assess gephyrin PTMs and identify potential additional therapeutic targets. This knowledge will facilitate the design of procedures to moderate BZD tolerance and improve future GABA_A_R-targeted drug development.

### Limitations of the Study

There were several limitations in this study. Firstly, the findings could be strengthened by the addition of experiments in which co-treatment with the BZD-site antagonist flumazenil is used to confirm BZD-specific effects. Secondly, DNA-PAINT experiments would be improved by three-dimensional imaging to simultaneously visualize presynaptic active-zone proteins such as RIM, as the current data do not provide insight into trans-synaptic alignment of SSDs after chronic BZD treatment. Finally, while neuronal cell culture was used here to examine molecular mechanisms with high resolution, future studies will be needed to determine the relevance of these findings *in vivo* and assess potential sex-specific differences. This is particularly true given that our prior studies of 7-day DZP treatment in mice showed increased γ2-GABA_A_R synaptic expression and increased extrasynaptic gephyrin expression without loss of synaptic or total scaffold (Lorenz-Guertin et al., 2023). Differential timelines of neuroadaptations *in vitro* versus *in vivo* are a potential contributing factor, which could be elucidated in future research through the analysis of additional time points of BZD treatment in both systems.

## Author contributions

**Caitlyn A. Chapman:** Conceptualization, Methodology, Validation, Formal analysis, Investigation, Writing – Original Draft, Writing – Review & Editing, Visualization; performed immunofluorescence, DNA-PAINT, biochemical, and live-imaging experiments and analysis and performed imaging and analysis of proximity ligation assay experiments. **Nadya Povysheva:** Methodology, Validation, Formal analysis, Investigation, Writing – Review & Editing; performed electrophysiology experiments and analysis. **Tyler Tarr:** Writing - Review & Editing, Validation, Methodology, Formal analysis, Visualization, Software; wrote and edited MATLAB scripts for DNA-PAINT experiments and assisted with analysis. **Jessica L. Nuwer:** Methodology, Validation, Investigation, Writing – Review & Editing; performed proximity ligation assay experiments. **Stephen D. Meriney:** Writing – Review & Editing, Funding acquisition. **Jon W. Johnson:** Writing – Review & Editing, Funding acquisition**. Tija C. Jacob:** Conceptualization, Methodology, Writing – Review & Editing, Supervision, Project administration, Funding acquisition.

## Data availability

Data and MATLAB codes will be supplied upon request.

## Acknowledgments

Work performed in the University of Pittsburgh Dietrich School Microscopy and Imaging Suite (RRID:SCR_022084) and services and instruments used in this project were graciously supported, in part, by the University of Pittsburgh. Graphical abstract and schematics were created in https://BioRender.com.

## References

Alldred, M. J., Mulder-Rosi, J., Lingenfelter, S. E., Chen, G., and Lüscher, B. (2005). Distinct gamma2 subunit domains mediate clustering and synaptic function of postsynaptic GABAA receptors and gephyrin. J. Neurosci. 25, 594–603. doi:10.1523/JNEUROSCI.4011-04.2005.

Allison, C., and Pratt, J. A. (2006). Differential effects of two chronic diazepam treatment regimes on withdrawal anxiety and AMPA receptor characteristics. Neuropsychopharmacology 31, 602–619. doi:10.1038/sj.npp.1300800.

Anderson, M. C., Levy, A. D., Dharmasri, P. A., Metzbower, S. R., and Blanpied, T. A. (2023). Trans-synaptic molecular context of NMDA receptor nanodomains. BioRxiv. doi:10.1101/2023.12.22.573055.

Arnot, M. I., Davies, M., Martin, I. L., and Bateson, A. N. (2001). GABA(A) receptor gene expression in rat cortex: differential effects of two chronic diazepam treatment regimes. J. Neurosci. Res. 64, 617–625. doi:10.1002/jnr.1115.

Bachhuber, M. A., Hennessy, S., Cunningham, C. O., and Starrels, J. L. (2016). Increasing benzodiazepine prescriptions and overdose mortality in the United States, 1996-2013. Am. J. Public Health 106, 686–688. doi:10.2105/AJPH.2016.303061.

Barberis, A. (2020). Postsynaptic plasticity of GABAergic synapses. Neuropharmacology 169, 107643. doi:10.1016/j.neuropharm.2019.05.020.

Bateson, A. N. (2002). Basic pharmacologic mechanisms involved in benzodiazepine tolerance and withdrawal. Curr. Pharm. Des. 8, 5–21. doi:10.2174/1381612023396681.

Battaglia, S., Renner, M., Russeau, M., Côme, E., Tyagarajan, S. K., and Lévi, S. (2018). Activity-Dependent Inhibitory Synapse Scaling Is Determined by Gephyrin Phosphorylation and Subsequent Regulation of GABAA Receptor Diffusion. eNeuro 5. doi:10.1523/ENEURO.0203-17.2017.

Bensussen, S., Shankar, S., Ching, K. H., Zemel, D., Ta, T. L., Mount, R. A., Shroff, S. N., Gritton, H. J., Fabris, P., Vanbenschoten, H., et al. (2020). A viral toolbox of genetically encoded fluorescent synaptic tags. iScience 23, 101330. doi:10.1016/j.isci.2020.101330.

Bevers, M. B., and Neumar, R. W. (2008). Mechanistic role of calpains in postischemic neurodegeneration. J. Cereb. Blood Flow Metab. 28, 655–673. doi:10.1038/sj.jcbfm.9600595.

Bogdanov, Y., Michels, G., Armstrong-Gold, C., Haydon, P. G., Lindstrom, J., Pangalos, M., and Moss, S. J. (2006). Synaptic GABAA receptors are directly recruited from their extrasynaptic counterparts. EMBO J. 25, 4381–4389. doi:10.1038/sj.emboj.7601309.

Brady, M. L., and Jacob, T. C. (2015). Synaptic localization of α5 GABA (A) receptors via gephyrin interaction regulates dendritic outgrowth and spine maturation. Dev. Neurobiol. 75, 1241–1251. doi:10.1002/dneu.22280.

Carricaburu, E., Benner, O., Burlingham, S. R., Dos Santos Passos, C., Hobaugh, N., Karr, C. H., and Chanda, S. (2024). Gephyrin promotes autonomous assembly and synaptic localization of GABAergic postsynaptic components without presynaptic GABA release. Proc Natl Acad Sci USA 121, e2315100121. doi:10.1073/pnas.2315100121.

Chapoutot, M., Peter-Derex, L., Bastuji, H., Leslie, W., Schoendorff, B., Heinzer, R., Siclari, F., Nicolas, A., Lemoine, P., Higgins, S., et al. (2021). Cognitive Behavioral Therapy and Acceptance and Commitment Therapy for the Discontinuation of Long-Term Benzodiazepine Use in Insomnia and Anxiety Disorders. Int. J. Environ. Res. Public Health 18. doi:10.3390/ijerph181910222.

Chen, H., Tang, A.-H., and Blanpied, T. A. (2018). Subsynaptic spatial organization as a regulator of synaptic strength and plasticity. Curr. Opin. Neurobiol. 51, 147–153. doi:10.1016/j.conb.2018.05.004.

Chen, J.-H., Blanpied, T. A., and Tang, A.-H. (2020). Quantification of trans-synaptic protein alignment: A data analysis case for single-molecule localization microscopy. Methods 174, 72–80. doi:10.1016/j.ymeth.2019.07.016.

Choii, G., and Ko, J. (2015). Gephyrin: a central GABAergic synapse organizer. Exp. Mol. Med. 47, e158. doi:10.1038/emm.2015.5.

Choquet, D., and Triller, A. (2013). The dynamic synapse. Neuron 80, 691–703. doi:10.1016/j.neuron.2013.10.013.

Costa, J. T., Mele, M., Baptista, M. S., Gomes, J. R., Ruscher, K., Nobre, R. J., de Almeida, L. P., Wieloch, T., and Duarte, C. B. (2016). Gephyrin cleavage in in vitro brain ischemia decreases GABAA receptor clustering and contributes to neuronal death. Mol. Neurobiol. 53, 3513–3527. doi:10.1007/s12035-015-9283-2.

Crosby, K. C., Gookin, S. E., Garcia, J. D., Hahm, K. M., Dell’Acqua, M. L., and Smith, K. R. (2019). Nanoscale subsynaptic domains underlie the organization of the inhibitory synapse. Cell Rep. 26, 3284–3297.e3. doi:10.1016/j.celrep.2019.02.070.

Danglot, L., Triller, A., and Bessis, A. (2003). Association of gephyrin with synaptic and extrasynaptic GABAA receptors varies during development in cultured hippocampal neurons. Mol. Cell. Neurosci. 23, 264–278. doi:10.1016/s1044-7431(03)00069-1.

Dani, A., Huang, B., Bergan, J., Dulac, C., and Zhuang, X. (2010). Superresolution imaging of chemical synapses in the brain. Neuron 68, 843–856. doi:10.1016/j.neuron.2010.11.021.

Davenport, E. C., Pendolino, V., Kontou, G., McGee, T. P., Sheehan, D. F., López-Doménech, G., Farrant, M., and Kittler, J. T. (2017). An Essential Role for the Tetraspanin LHFPL4 in the Cell-Type-Specific Targeting and Clustering of Synaptic GABAA Receptors. Cell Rep. 21, 70–83. doi:10.1016/j.celrep.2017.09.025.

Dejanovic, B., and Schwarz, G. (2014). Neuronal nitric oxide synthase-dependent S-nitrosylation of gephyrin regulates gephyrin clustering at GABAergic synapses. J. Neurosci. 34, 7763–7768. doi:10.1523/JNEUROSCI.0531-14.2014.

Dejanovic, B., Semtner, M., Ebert, S., Lamkemeyer, T., Neuser, F., Lüscher, B., Meier, J. C., and Schwarz, G. (2014). Palmitoylation of gephyrin controls receptor clustering and plasticity of GABAergic synapses. PLoS Biol. 12, e1001908. doi:10.1371/journal.pbio.1001908.

Dharmasri, P. A., DeMarco, E. M., Anderson, M. C., Levy, A. D., and Blanpied, T. A. (2024). Loss of postsynaptic NMDARs drives nanoscale reorganization of Munc13-1 and PSD-95. *BioRxiv*. doi:10.1101/2024.01.12.574705.

Ehrensperger, M.-V., Hanus, C., Vannier, C., Triller, A., and Dahan, M. (2007). Multiple association states between glycine receptors and gephyrin identified by SPT analysis. Biophys. J. 92, 3706–3718. doi:10.1529/biophysj.106.095596.

Essrich, C., Lorez, M., Benson, J. A., Fritschy, J. M., and Lüscher, B. (1998). Postsynaptic clustering of major GABAA receptor subtypes requires the gamma 2 subunit and gephyrin. Nat. Neurosci. 1, 563–571. doi:10.1038/2798.

Fernandes, C., and File, S. E. (1999). Dizocilpine does not prevent the development of tolerance to the anxiolytic effects of diazepam in rats. Brain Res. 815, 431–434. doi:10.1016/s0006-8993(98)01160-3.

Ferreri, M. C., Gutiérrez, M. L., and Gravielle, M. C. (2015). Tolerance to the sedative and anxiolytic effects of diazepam is associated with different alterations of GABAA receptors in rat cerebral cortex. Neuroscience 310, 152–162. doi:10.1016/j.neuroscience.2015.09.038.

Flores, C. E., Nikonenko, I., Mendez, P., Fritschy, J.-M., Tyagarajan, S. K., and Muller, D. (2015). Activity-dependent inhibitory synapse remodeling through gephyrin phosphorylation. Proc Natl Acad Sci USA 112, E65–72. doi:10.1073/pnas.1411170112.

Foitzick, M. F., Medina, N. B., Iglesias García, L. C., and Gravielle, M. C. (2020). Benzodiazepine exposure induces transcriptional down-regulation of GABAA receptor α1 subunit gene via L-type voltage-gated calcium channel activation in rat cerebrocortical neurons. Neurosci. Lett. 721, 134801. doi:10.1016/j.neulet.2020.134801.

Furukawa, T., Shimoyama, S., Miki, Y., Nikaido, Y., Koga, K., Nakamura, K., Wakabayashi, K., and Ueno, S. (2017). Chronic diazepam administration increases the expression of Lcn2 in the CNS. Pharmacol. Res. Perspect. 5, e00283. doi:10.1002/prp2.283.

Gao, L., and Greenfield, L. J. (2005). Activation of protein kinase C reduces benzodiazepine potency at GABAA receptors in NT2-N neurons. Neuropharmacology 48, 333–342. doi:10.1016/j.neuropharm.2004.10.010.

Garcia, J. D., Gookin, S. E., Crosby, K. C., Schwartz, S. L., Tiemeier, E., Kennedy, M. J., Dell’Acqua, M. L., Herson, P. S., Quillinan, N., and Smith, K. R. (2021). Stepwise disassembly of GABAergic synapses during pathogenic excitotoxicity. Cell Rep. 37, 110142. doi:10.1016/j.celrep.2021.110142.

Gerlach, L. B., Strominger, J., Kim, H. M., and Maust, D. T. (2019). Discontinuation of chronic benzodiazepine use among adults in the united states. J. Gen. Intern. Med. 34, 1833–1840. doi:10.1007/s11606-019-05098-0.

Ghosh, H., Auguadri, L., Battaglia, S., Simone Thirouin, Z., Zemoura, K., Messner, S., Acuña, M. A., Wildner, H., Yévenes, G. E., Dieter, A., et al. (2016). Several posttranslational modifications act in concert to regulate gephyrin scaffolding and GABAergic transmission. Nat. Commun. 7, 13365. doi:10.1038/ncomms13365.

Gielen, M. C., Lumb, M. J., and Smart, T. G. (2012). Benzodiazepines modulate GABAA receptors by regulating the preactivation step after GABA binding. J. Neurosci. 32, 5707–5715. doi:10.1523/JNEUROSCI.5663-11.2012.

Goebel-Goody, S. M., Davies, K. D., Alvestad Linger, R. M., Freund, R. K., and Browning, M. D. (2009). Phospho-regulation of synaptic and extrasynaptic N-methyl-d-aspartate receptors in adult hippocampal slices. Neuroscience 158, 1446–1459. doi:10.1016/j.neuroscience.2008.11.006.

González Gómez, L. C., Medina, N. B., Sanz Blasco, S., and Gravielle, M. C. (2023). Diazepam-Induced Down-Regulation of the GABAA receptor α1 Subunit, as mediated by the activation of L-Type Voltage-Gated calcium Channel/Ca2+/Protein kinase a signaling cascade. Neurosci. Lett. 810, 137358. doi:10.1016/j.neulet.2023.137358.

Gookin, S. E., Taylor, M. R., Schwartz, S. L., Kennedy, M. J., Dell’Acqua, M. L., Crosby, K. C., and Smith, K. R. (2022). Complementary Use of Super-Resolution Imaging Modalities to Study the Nanoscale Architecture of Inhibitory Synapses. Front. Synaptic Neurosci. 14, 852227. doi:10.3389/fnsyn.2022.852227.

Gouzer, G., Specht, C. G., Allain, L., Shinoe, T., and Triller, A. (2014). Benzodiazepine-dependent stabilization of GABA(A) receptors at synapses. Mol. Cell. Neurosci. 63, 101–113. doi:10.1016/j.mcn.2014.10.004.

Groeneweg, F. L., Trattnig, C., Kuhse, J., Nawrotzki, R. A., and Kirsch, J. (2018). Gephyrin: a key regulatory protein of inhibitory synapses and beyond. Histochem. Cell Biol. 150, 489–508. doi:10.1007/s00418-018-1725-2.

Gross, G. G., Junge, J. A., Mora, R. J., Kwon, H.-B., Olson, C. A., Takahashi, T. T., Liman, E. R., Ellis-Davies, G. C. R., McGee, A. W., Sabatini, B. L., et al. (2013). Recombinant probes for visualizing endogenous synaptic proteins in living neurons. Neuron 78, 971–985. doi:10.1016/j.neuron.2013.04.017.

Günther, U., Benson, J., Benke, D., Fritschy, J. M., Reyes, G., Knoflach, F., Crestani, F., Aguzzi, A., Arigoni, M., Lang, Y., et al. (1995). Benzodiazepine-insensitive mice generated by targeted disruption of the gamma 2 subunit gene of gamma-aminobutyric acid type A receptors. Proc Natl Acad Sci USA 92, 7749–7753. doi:10.1073/pnas.92.17.7749.

Holt, R. A., Bateson, A. N., and Martin, I. L. (1996). Chronic treatment with diazepam or abecarnil differently affects the expression of GABAA receptor subunit mRNAs in the rat cortex. Neuropharmacology 35, 1457– 1463. doi:10.1016/s0028-3908(96)00064-0.

Hu, X. J., and Ticku, M. K. (1994). Chronic benzodiazepine agonist treatment produces functional uncoupling of the gamma-aminobutyric acid-benzodiazepine receptor ionophore complex in cortical neurons. Mol. Pharmacol. 45, 618–625.

Impagnatiello, F., Pesold, C., Longone, P., Caruncho, H., Fritschy, J. M., Costa, E., and Guidotti, A. (1996). Modifications of gamma-aminobutyric acidA receptor subunit expression in rat neocortex during tolerance to diazepam. Mol. Pharmacol. 49, 822–831.

Jacob, T. C., Bogdanov, Y. D., Magnus, C., Saliba, R. S., Kittler, J. T., Haydon, P. G., and Moss, S. J. (2005). Gephyrin regulates the cell surface dynamics of synaptic GABAA receptors. J. Neurosci. 25, 10469–10478. doi:10.1523/JNEUROSCI.2267-05.2005.

Jacob, T. C., Michels, G., Silayeva, L., Haydon, J., Succol, F., and Moss, S. J. (2012). Benzodiazepine treatment induces subtype-specific changes in GABA(A) receptor trafficking and decreases synaptic inhibition. Proc Natl Acad Sci USA 109, 18595–18600. doi:10.1073/pnas.1204994109.

Jacob, T. C., Moss, S. J., and Jurd, R. (2008). GABA(A) receptor trafficking and its role in the dynamic modulation of neuronal inhibition. Nat. Rev. Neurosci. 9, 331–343. doi:10.1038/nrn2370.

Janhsen, K., Roser, P., and Hoffmann, K. (2015). The problems of long-term treatment with benzodiazepines and related substances. Dtsch Arztebl Int 112, 1–7. doi:10.3238/arztebl.2015.0001.

Jungmann, R., Steinhauer, C., Scheible, M., Kuzyk, A., Tinnefeld, P., and Simmel, F. C. (2010). Single-molecule kinetics and super-resolution microscopy by fluorescence imaging of transient binding on DNA origami. Nano Lett. 10, 4756–4761. doi:10.1021/nl103427w.

Kasaragod, V. B., and Schindelin, H. (2018). Structure-Function Relationships of Glycine and GABAA Receptors and Their Interplay With the Scaffolding Protein Gephyrin. Front. Mol. Neurosci. 11, 317. doi:10.3389/fnmol.2018.00317.

Kaufmann, C. N., Spira, A. P., Depp, C. A., and Mojtabai, R. (2018). Long-Term Use of Benzodiazepines and Nonbenzodiazepine Hypnotics, 1999-2014. Psychiatr. Serv. 69, 235–238. doi:10.1176/appi.ps.201700095.

Kawasaki, B. T., Hoffman, K. B., Yamamoto, R. S., and Bahr, B. A. (1997). Variants of the receptor/channel clustering molecule gephyrin in brain: Distinct distribution patterns, developmental profiles, and proteolytic cleavage by calpain. Journal of Neuroscience Research.

Kneussel, M., Brandstätter, J. H., Laube, B., Stahl, S., Müller, U., and Betz, H. (1999). Loss of postsynaptic GABA(A) receptor clustering in gephyrin-deficient mice. J. Neurosci. 19, 9289–9297. doi:10.1523/JNEUROSCI.19-21-09289.1999.

Kowalczyk, S., Winkelmann, A., Smolinsky, B., Förstera, B., Neundorf, I., Schwarz, G., and Meier, J. C. (2013). Direct binding of GABAA receptor β2 and β3 subunits to gephyrin. Eur. J. Neurosci. 37, 544–554. doi:10.1111/ejn.12078.

Kurko, T. A. T., Saastamoinen, L. K., Tähkäpää, S., Tuulio-Henriksson, A., Taiminen, T., Tiihonen, J., Airaksinen, M. S., and Hietala, J. (2015). Long-term use of benzodiazepines: Definitions, prevalence and usage patterns - a systematic review of register-based studies. Eur. Psychiatry 30, 1037–1047. doi:10.1016/j.eurpsy.2015.09.003.

Lévi, S., Le Roux, N., Eugène, E., and Poncer, J. C. (2015). Benzodiazepine ligands rapidly influence GABAA receptor diffusion and clustering at hippocampal inhibitory synapses. Neuropharmacology 88, 199–208. doi:10.1016/j.neuropharm.2014.06.002.

Li, M., Szabo, A., and Rosenberg, H. C. (2000). Down-regulation of benzodiazepine binding to alpha 5 subunit-containing gamma-aminobutyric Acid(A) receptors in tolerant rat brain indicates particular involvement of the hippocampal CA1 region. J. Pharmacol. Exp. Ther. 295, 689–696.

Longone, P., Impagnatiello, F., Guidotti, A., and Costa, E. (1996). Reversible Modification of GABA A Receptor Subunit mRNA Expression During Tolerance to Diazepam-induced Cognition Dysfunction. Neuropharmacology 35, 1465–1473. doi:10.1016/S0028-3908(96)00071-8.

Lorenz-Guertin, J. M., Bambino, M. J., Das, S., Weintraub, S. T., and Jacob, T. C. (2019). Diazepam accelerates GABAAR synaptic exchange and alters intracellular trafficking. Front. Cell. Neurosci. 13, 163. doi:10.3389/fncel.2019.00163.

Lorenz-Guertin, J. M., Povysheva, N., Chapman, C. A., MacDonald, M. L., Fazzari, M., Nigam, A., Nuwer, J. L., Das, S., Brady, M. L., Vajn, K., et al. (2023). Inhibitory and excitatory synaptic neuroadaptations in the diazepam tolerant brain. Neurobiol. Dis. 185, 106248. doi:10.1016/j.nbd.2023.106248.

Lorenz-Guertin, J. M., Wilcox, M. R., Zhang, M., Larsen, M. B., Pilli, J., Schmidt, B. F., Bruchez, M. P., Johnson, J. W., Waggoner, A. S., Watkins, S. C., et al. (2017). A versatile optical tool for studying synaptic GABAA receptor trafficking. J. Cell Sci. 130, 3933–3945. doi:10.1242/jcs.205286.

MacGillavry, H. D., Song, Y., Raghavachari, S., and Blanpied, T. A. (2013). Nanoscale scaffolding domains within the postsynaptic density concentrate synaptic AMPA receptors. Neuron 78, 615–622. doi:10.1016/j.neuron.2013.03.009.

Malherbe, P., Sigel, E., Baur, R., Persohn, E., Richards, J. G., and Mohler, H. (1990). Functional characteristics and sites of gene expression of the alpha 1, beta 1, gamma 2-isoform of the rat GABAA receptor. J. Neurosci. 10, 2330–2337. doi:10.1523/JNEUROSCI.10-07-02330.1990.

Martenson, J. S., Yamasaki, T., Chaudhury, N. H., Albrecht, D., and Tomita, S. (2017). Assembly rules for GABAA receptor complexes in the brain. eLife 6. doi:10.7554/eLife.27443.

Maust, D. T., Lin, L. A., and Blow, F. C. (2019). Benzodiazepine use and misuse among adults in the United States. Psychiatr. Serv. 70, 97–106. doi:10.1176/appi.ps.201800321.

Mele, M., Leal, G., and Duarte, C. B. (2016). Role of GABAA R trafficking in the plasticity of inhibitory synapses. J. Neurochem. 139, 997–1018. doi:10.1111/jnc.13742.

Merlaud, Z., Marques, X., Russeau, M., Saade, U., Tostain, M., Moutkine, I., Gielen, M., Corringer, P.-J., and Lévi, S. (2022). Conformational state-dependent regulation of GABAA receptor diffusion and subsynaptic domains. iScience 25, 105467. doi:10.1016/j.isci.2022.105467.

Morin, C. M., Bélanger, L., Bastien, C., and Vallières, A. (2005). Long-term outcome after discontinuation of benzodiazepines for insomnia: a survival analysis of relapse. Behav. Res. Ther. 43, 1–14. doi:10.1016/j.brat.2003.12.002.

Mozrzymas, J. W., Wójtowicz, T., Piast, M., Lebida, K., Wyrembek, P., and Mercik, K. (2007). GABA transient sets the susceptibility of mIPSCs to modulation by benzodiazepine receptor agonists in rat hippocampal neurons. J Physiol (Lond) 585, 29–46. doi:10.1113/jphysiol.2007.143602.

Mukherjee, J., Kretschmannova, K., Gouzer, G., Maric, H.-M., Ramsden, S., Tretter, V., Harvey, K., Davies, P. A., Triller, A., Schindelin, H., et al. (2011). The residence time of GABA(A)Rs at inhibitory synapses is determined by direct binding of the receptor α1 subunit to gephyrin. J. Neurosci. 31, 14677–14687. doi:10.1523/JNEUROSCI.2001-11.2011.

Nair, D., Hosy, E., Petersen, J. D., Constals, A., Giannone, G., Choquet, D., and Sibarita, J.-B. (2013). Super-resolution imaging reveals that AMPA receptors inside synapses are dynamically organized in nanodomains regulated by PSD95. J. Neurosci. 33, 13204–13224. doi:10.1523/JNEUROSCI.2381-12.2013.

Nicholson, M. W., Sweeney, A., Pekle, E., Alam, S., Ali, A. B., Duchen, M., and Jovanovic, J. N. (2018). Diazepam-induced loss of inhibitory synapses mediated by PLCδ/ Ca2+/calcineurin signalling downstream of GABAA receptors. Mol. Psychiatry 23, 1851–1867. doi:10.1038/s41380-018-0100-y.

Niwa, F., Bannai, H., Arizono, M., Fukatsu, K., Triller, A., and Mikoshiba, K. (2012). Gephyrin-independent GABA(A)R mobility and clustering during plasticity. PLoS ONE 7, e36148. doi:10.1371/journal.pone.0036148.

Nuwer, J. L., Brady, M. L., Povysheva, N. V., Coyne, A., and Jacob, T. C. (2021). Sustained treatment with an α5 GABA A receptor negative allosteric modulator delays excitatory circuit development while maintaining GABAergic neurotransmission. Neuropharmacology 197, 108724. doi:10.1016/j.neuropharm.2021.108724.

Nuwer, J. L., Povysheva, N., and Jacob, T. C. (2023). Long-term α5 GABA A receptor negative allosteric modulator treatment reduces NMDAR-mediated neuronal excitation and maintains basal neuronal inhibition. Neuropharmacology 237, 109587. doi:10.1016/j.neuropharm.2023.109587.

Olah, S. S., Kareemo, D. J., Buchta, W. C., Sinnen, B. L., Miller, C. N., Actor-Engel, H. S., Gookin, S. E., Winborn, C. S., Kleinjan, M. S., Crosby, K. C., et al. (2023). Acute reorganization of postsynaptic GABAA receptors reveals the functional impact of molecular nanoarchitecture at inhibitory synapses. Cell Rep. 42, 113331. doi:10.1016/j.celrep.2023.113331.

Olfson, M., King, M., and Schoenbaum, M. (2015). Benzodiazepine use in the United States. JAMA Psychiatry 72, 136–142. doi:10.1001/jamapsychiatry.2014.1763.

Olsen, R. W., and Sieghart, W. (2009). GABA A receptors: subtypes provide diversity of function and pharmacology. Neuropharmacology 56, 141–148. doi:10.1016/j.neuropharm.2008.07.045.

Olsen, R. W., and Sieghart, W. (2008). International Union of Pharmacology. LXX. Subtypes of gamma-aminobutyric acid(A) receptors: classification on the basis of subunit composition, pharmacology, and function. Update. Pharmacol. Rev. 60, 243–260. doi:10.1124/pr.108.00505.

Pennacchietti, F., Vascon, S., Nieus, T., Rosillo, C., Das, S., Tyagarajan, S. K., Diaspro, A., Del Bue, A., Petrini, E. M., Barberis, A., et al. (2017). Nanoscale molecular reorganization of the inhibitory postsynaptic density is a determinant of gabaergic synaptic potentiation. J. Neurosci. 37, 1747–1756. doi:10.1523/JNEUROSCI.0514-16.2016.

Pesold, C., Caruncho, H. J., Impagnatiello, F., Berg, M. J., Fritschy, J. M., Guidotti, A., and Costa, E. (1997). Tolerance to diazepam and changes in GABAA receptor subunit expression in rat neocortical areas. Neuroscience 79, 477–487. doi:10.1016/S0306-4522(96)00609-4.

Petrini, E. M., and Barberis, A. (2014). Diffusion dynamics of synaptic molecules during inhibitory postsynaptic plasticity. Front. Cell. Neurosci. 8, 300. doi:10.3389/fncel.2014.00300.

Petrini, E. M., Ravasenga, T., Hausrat, T. J., Iurilli, G., Olcese, U., Racine, V., Sibarita, J.-B., Jacob, T. C., Moss, S. J., Benfenati, F., et al. (2014). Synaptic recruitment of gephyrin regulates surface GABAA receptor dynamics for the expression of inhibitory LTP. Nat. Commun. 5, 3921. doi:10.1038/ncomms4921.

Pétursson, H. (1994). The benzodiazepine withdrawal syndrome. Addiction 89, 1455–1459. doi:10.1111/j.1360-0443.1994.tb03743.x.

Pizzarelli, R., Griguoli, M., Zacchi, P., Petrini, E. M., Barberis, A., Cattaneo, A., and Cherubini, E. (2020). Tuning gabaergic inhibition: gephyrin molecular organization and functions. Neuroscience 439, 125–136. doi:10.1016/j.neuroscience.2019.07.036.

Povysheva, N. V., and Johnson, J. W. (2016). Effects of memantine on the excitation-inhibition balance in prefrontal cortex. Neurobiol. Dis. 96, 75–83. doi:10.1016/j.nbd.2016.08.006.

Pritchett, D. B., Sontheimer, H., Shivers, B. D., Ymer, S., Kettenmann, H., Schofield, P. R., and Seeburg, P. H. (1989). Importance of a novel GABAA receptor subunit for benzodiazepine pharmacology. Nature 338, 582–585. doi:10.1038/338582a0.

Renner, M., Schweizer, C., Bannai, H., Triller, A., and Lévi, S. (2012). Diffusion barriers constrain receptors at synapses. PLoS ONE 7, e43032. doi:10.1371/journal.pone.0043032.

Sahu, M. P., Nikkilä, O., Lågas, S., Kolehmainen, S., and Castrén, E. (2019). Culturing primary neurons from rat hippocampus and cortex. Neuronal Signal. 3, NS20180207. doi:10.1042/NS20180207.

Sander, B., Tria, G., Shkumatov, A. V., Kim, E.-Y., Grossmann, J. G., Tessmer, I., Svergun, D. I., and Schindelin, H. (2013). Structural characterization of gephyrin by AFM and SAXS reveals a mixture of compact and extended states. Acta Crystallogr. D Biol. Crystallogr. 69, 2050–2060. doi:10.1107/S0907444913018714.

Schnitzbauer, J., Strauss, M. T., Schlichthaerle, T., Schueder, F., and Jungmann, R. (2017). Super-resolution microscopy with DNA-PAINT. Nat. Protoc. 12, 1198–1228. doi:10.1038/nprot.2017.024.

Schweizer, C. (2003). The γ2 subunit of GABAA receptors is required for maintenance of receptors at mature synapses. Molecular and Cellular Neuroscience 24, 442–450. doi:10.1016/S1044-7431(03)00202-1.

Sharma, D., Dixit, A. B., Dey, S., Tripathi, M., Doddamani, R., Sharma, M. C., Lalwani, S., Gurjar, H. K., Chandra, P. S., and Banerjee, J. (2021). Increased levels of α4-containing GABAA receptors in focal cortical dysplasia: A possible cause of benzodiazepine resistance. Neurochem. Int. 148, 105084. doi:10.1016/j.neuint.2021.105084.

Shen, Z.-C., Wu, P.-F., Wang, F., Xia, Z.-X., Deng, Q., Nie, T.-L., Zhang, S.-Q., Zheng, H.-L., Liu, W.-H., Lu, J.-J., et al. (2019). Gephyrin palmitoylation in basolateral amygdala mediates the anxiolytic action of benzodiazepine. Biol. Psychiatry 85, 202–213. doi:10.1016/j.biopsych.2018.09.024.

Sograte-Idrissi, S., Schlichthaerle, T., Duque-Afonso, C. J., Alevra, M., Strauss, S., Moser, T., Jungmann, R., Rizzoli, S. O., and Opazo, F. (2020). Circumvention of common labelling artefacts using secondary nanobodies. Nanoscale 12, 10226–10239. doi:10.1039/d0nr00227e.

Specht, C. G., Izeddin, I., Rodriguez, P. C., El Beheiry, M., Rostaing, P., Darzacq, X., Dahan, M., and Triller, A. (2013). Quantitative nanoscopy of inhibitory synapses: counting gephyrin molecules and receptor binding sites. Neuron 79, 308–321. doi:10.1016/j.neuron.2013.05.013.

Talos, D. M., Sun, H., Kosaras, B., Joseph, A., Folkerth, R. D., Poduri, A., Madsen, J. R., Black, P. M., and Jensen, F. E. (2012). Altered inhibition in tuberous sclerosis and type IIb cortical dysplasia. Ann. Neurol. 71, 539–551. doi:10.1002/ana.22696.

Tang, A.-H., Chen, H., Li, T. P., Metzbower, S. R., MacGillavry, H. D., and Blanpied, T. A. (2016). A trans-synaptic nanocolumn aligns neurotransmitter release to receptors. Nature 536, 210–214. doi:10.1038/nature19058.

Tanguay Bernard, M.-M., Luc, M., Carrier, J.-D., Fournier, L., Duhoux, A., Côté, E., Lessard, O., Gibeault, C., Bocti, C., and Roberge, P. (2018). Patterns of benzodiazepines use in primary care adults with anxiety disorders. Heliyon 4, e00688. doi:10.1016/j.heliyon.2018.e00688.

Tretter, V., Jacob, T. C., Mukherjee, J., Fritschy, J.-M., Pangalos, M. N., and Moss, S. J. (2008). The clustering of GABA(A) receptor subtypes at inhibitory synapses is facilitated via the direct binding of receptor alpha 2 subunits to gephyrin. J. Neurosci. 28, 1356–1365. doi:10.1523/JNEUROSCI.5050-07.2008.

Tretter, V., Kerschner, B., Milenkovic, I., Ramsden, S. L., Ramerstorfer, J., Saiepour, L., Maric, H.-M., Moss, S. J., Schindelin, H., Harvey, R. J., et al. (2011). Molecular basis of the γ-aminobutyric acid A receptor α3 subunit interaction with the clustering protein gephyrin. J. Biol. Chem. 286, 37702–37711. doi:10.1074/jbc.M111.291336.

Tyagarajan, S. K., Ghosh, H., Yévenes, G. E., Imanishi, S. Y., Zeilhofer, H. U., Gerrits, B., and Fritschy, J.-M. (2013). Extracellular signal-regulated kinase and glycogen synthase kinase 3β regulate gephyrin postsynaptic aggregation and GABAergic synaptic function in a calpain-dependent mechanism. J. Biol. Chem. 288, 9634–9647. doi:10.1074/jbc.M112.442616.

Tyagarajan, S. K., Ghosh, H., Yévenes, G. E., Nikonenko, I., Ebeling, C., Schwerdel, C., Sidler, C., Zeilhofer, H. U., Gerrits, B., Muller, D., et al. (2011). Regulation of GABAergic synapse formation and plasticity by GSK3beta-dependent phosphorylation of gephyrin. Proc Natl Acad Sci USA 108, 379–384. doi:10.1073/pnas.1011824108.

Vinkers, C. H., and Olivier, B. (2012). Mechanisms Underlying Tolerance after Long-Term Benzodiazepine Use: A Future for Subtype-Selective GABA(A) Receptor Modulators? Adv. Pharmacol. Sci. 2012, 416864. doi:10.1155/2012/416864.

Vlachos, A., Reddy-Alla, S., Papadopoulos, T., Deller, T., and Betz, H. (2013). Homeostatic regulation of gephyrin scaffolds and synaptic strength at mature hippocampal GABAergic postsynapses. Cereb. Cortex 23, 2700–2711. doi:10.1093/cercor/bhs260.

Vosler, P. S., Brennan, C. S., and Chen, J. (2008). Calpain-mediated signaling mechanisms in neuronal injury and neurodegeneration. Mol. Neurobiol. 38, 78–100. doi:10.1007/s12035-008-8036-x.

Weibrecht, I., Leuchowius, K.-J., Clausson, C.-M., Conze, T., Jarvius, M., Howell, W. M., Kamali-Moghaddam, M., and Söderberg, O. (2010). Proximity ligation assays: a recent addition to the proteomics toolbox. Expert Rev. Proteomics 7, 401–409. doi:10.1586/epr.10.10.

Werner, C., Sauer, M., and Geis, C. (2021). Super-resolving Microscopy in Neuroscience. Chem. Rev. 121, 11971–12015. doi:10.1021/acs.chemrev.0c01174.

Wright, B. T., Gluszek, C. F., and Heldt, S. A. (2014). The effects of repeated zolpidem treatment on tolerance, withdrawal-like symptoms, and GABAA receptor mRNAs profile expression in mice: comparison with diazepam. Psychopharmacology (Berl) 231, 2967–2979. doi:10.1007/s00213-014-3473-x.

Wu, Y., Rosenberg, H. C., Chiu, T. H., and Zhao, T. J. (1994). Subunit- and brain region-specific reduction of GABAA receptor subunit mRNAs during chronic treatment of rats with diazepam. J. Mol. Neurosci. 5, 105–120. doi:10.1007/BF02736752.

Yamasaki, T., Hoyos-Ramirez, E., Martenson, J. S., Morimoto-Tomita, M., and Tomita, S. (2017). GARLH family proteins stabilize GABAA receptors at synapses. Neuron 93, 1138–1152.e6. doi:10.1016/j.neuron.2017.02.023.

Yang, P.-F., Nie, T.-L., Sun, X.-N., Xu, L.-X., Ma, C., Wang, F., Long, L.-H., and Chen, J.-G. (2024). Wheel-Running Exercise Alleviates Anxiety-Like Behavior via Down-Regulating S-Nitrosylation of Gephyrin in the Basolateral Amygdala of Male Rats. Adv Sci (Weinh), e2400205. doi:10.1002/advs.202400205.

Yang, X., and Annaert, W. (2021). The nanoscopic organization of synapse structures: A common basis for cell communication. Membranes (Basel) 11. doi:10.3390/membranes11040248.

Yang, X., and Specht, C. G. (2019). Subsynaptic Domains in Super-Resolution Microscopy: The Treachery of Images. Front. Mol. Neurosci. 12, 161. doi:10.3389/fnmol.2019.00161.

Yan, Q., Schmidt, B. F., Perkins, L. A., Naganbabu, M., Saurabh, S., Andreko, S. K., and Bruchez, M. P. (2015). Near-instant surface-selective fluorogenic protein quantification using sulfonated triarylmethane dyes and fluorogen activating proteins. Org. Biomol. Chem. 13, 2078–2086. doi:10.1039/c4ob02309a.

Yu, W., Jiang, M., Miralles, C. P., Li, R.-W., Chen, G., and de Blas, A. L. (2007). Gephyrin clustering is required for the stability of GABAergic synapses. Mol. Cell. Neurosci. 36, 484–500. doi:10.1016/j.mcn.2007.08.008.

Zacchi, P., Antonelli, R., and Cherubini, E. (2014). Gephyrin phosphorylation in the functional organization and plasticity of GABAergic synapses. Front. Cell. Neurosci. 8, 103. doi:10.3389/fncel.2014.00103.

van Zundert, B., Castro, P., and Aguayo, L. G. (2005). Glycinergic and GABAergic synaptic transmission are differentially affected by gephyrin in spinal neurons. Brain Res. 1050, 40–47. doi:10.1016/j.brainres.2005.05.014.

