## Supplemental Table 1 for "Chronic benzodiazepine treatment triggers gephyrin scaffold destabilization and GABA_A_R subsynaptic reorganization"

### Supplementary Table S1: Summary of Statistics (Page 1 of 3)

| Location in Text: Section | Location in Text: Figure | Measurement | | Comparison | *N* Values (Independent Cultures) | *n* values (Veh, DZP) | Statistic (df) | Statistical test | *p* value | |
|---|---|---|---|---|---|---|---|---|---|---|
| Section 3.1 | 1B | DZP Potentiation of mIPSC Amplitude | | 7-day Veh + acute DZP | 3 | (5, 5) | t(4)=2.816 | paired *t*-test | ***0.0480*** | * |
| | | | | 7-day DZP + acute DZP | | | t(4)=2.278 | paired *t*-test | 0.0849 | ns |
| | 1C | % DZP Potentiation of mIPSC Amplitude | | 7-day Veh vs. DZP | | | t(8)=2.980 | unpaired *t*-test | ***0.0179*** | * |
| | 1D | DZP Potentiation of mIPSC Tau deceay | | 7-day Veh + acute DZP | | | t(4)=5.890 | paired *t*-test | ***0.0042*** | ** |
| | | | | 7-day DZP + acute DZP | | | t(4)=3.763 | paired *t*-test | ***0.0197*** | * |
| | 1E | % DZP Potentiation of mIPSC Tau decay | | 7-day Veh vs. DZP | | | t(8)=3.205 | unpaired *t*-test | ***0.0125*** | * |
| | S1B | Baseline mIPSC Amplitude | | 7-day Veh vs. DZP | 3 | (5, 5) | t(8)=1.649 | unpaired *t*-test | 0.1379 | ns |
| | | Baseline mIPSC Frequency | | | | | t(8)=0.6057 | unpaired *t*-test | 0.5615 | ns |
| | | Baseline mIPSC Tau decay | | | | | t(8)=1.541 | unpaired *t*-test | 0.1618 | ns |
| Section 3.2 | 3D | Total Synapse Area | γ2-GABA$_A$R | 7-day Veh vs DZP | 2 | Cells: (9, 7) Synapses: (216, 148) | - | Mann-Whitney test | ***0.005*** | *** |
| | | | gephyrin | | 2 | Cells: (9, 7) Synapses: (217, 148) | - | Mann-Whitney test | ***<0.0001*** | **** |
| | 3E | Total Synapse Localization Density | γ2-GABA$_A$R | 7-day Veh vs DZP | 2 | Cells: (9, 7) Synapses: (216, 148) | - | Mann-Whitney test | ***0.0019*** | ** |
| | | | gephyrin | | 2 | Cells: (9, 7) Synapses: (217, 148) | - | Mann-Whitney test | 0.1689 | ns |
| | 3G | SSDs per synapse | γ2-GABA$_A$R | 7-day Veh vs DZP | 2 | Cells: (9, 7) Synapses: (216, 148) | - | Mann-Whitney test | 0.2809 | ns |
| | | | gephyrin | | 2 | Cells: (9, 7) Synapses: (217, 148) | - | Mann-Whitney test | 0.1673 | ns |
| | 3H | SSD Area | γ2-GABA$_A$R | 7-day Veh vs DZP | 2 | Cells: (9, 7) SSD-containing synapses: (120, 84) | - | Mann-Whitney test | 0.2173 | ns |
| | | | gephyrin | | 2 | Cells: (9, 7) SSD-containing synapses: (213, 147) | - | Mann-Whitney test | ***0.0265*** | * |
| | 3I | SSD Localization Density | γ2-GABA$_A$R | 7-day Veh vs DZP | 2 | Cells: (9, 7) SSD-containing synapses: (120, 84) | - | Mann-Whitney test | 0.8527 | ns |
| | | | gephyrin | | 2 | Cells: (9, 7) SSD-containing synapses: (213, 147) | - | Mann-Whitney test | 0.3967 | ns |
| | 3J | Total SSD/Synapse Area | γ2-GABA$_A$R | 7-day Veh vs DZP | 2 | Cells: (9, 7) SSD-containing synapses: (120, 85) | - | Mann-Whitney test | 0.6347 | ns |
| | | | gephyrin | | 2 | Cells: (9, 7) SSD-containing synapses: (214, 148) | t(360)=0.7404 | unpaired t-test | 0.4595 | ns |
| | S2A | Enrichment Index: γ2-GABA$_A$R enriched to gephyrin | | 7-day Veh vs DZP | 2 | Cells: (9, 7) SSDs: (400, 267) | - | Mann-Whitney test | 0.2377 | ns |
| | S2B | Enrichment Index: gephyrin enriched to γ2-GABA$_A$R | | | 2 | Cells: (9, 7) SSDs: (188, 108) | - | Mann-Whitney test | 0.6056 | ns |
| Section 3.3 | 3C | PL Signal: Number of Objects | Whole-field mAb7a-gephyrin | 7-day Veh vs DZP | 3 | (37, 37) | - | Mann-Whitney test | ***0.0757*** | ns |
| | 3D | | Synaptic mAb7a-gephyrin | 7-day Veh vs DZP | | (36, 37) | - | Mann-Whitney test | ***0.0751*** | ns |
| | 3E | PL Signal: Sum Intensity | Whole-field mAb7a-gephyrin | 7-day Veh vs DZP | | (37, 37) | - | Mann-Whitney test | ***0.0877*** | ns |
| | 3F | | Synaptic mAb7a-gephyrin | 7-day Veh vs DZP | | (37, 36) | - | Mann-Whitney test | ***0.0263*** | * |

**Supplementary Table S1: Summary of Statistics (page 2 of 3)**

| Location in Text: Section | Location in Text: Figure | Measurement | | Comparison | *N* Values (Independent Cultures) | *n* values (Veh, DZP) | Statistic (df) | Statistical test | *p* value | |
|---|---|---|---|---|---|---|---|---|---|---|
| Section 3.3 | 4B | Full-length gephyrin | Synaptic | 7-day Veh vs DZP | 4 | (8, 8) | t(14)=2.084 | unpaired t-test | ***0.0559*** | ns |
| | | | Extrasynaptic | | | (8, 7) | t(14)=0.03822 | unpaired t-test | 0.1674 | ns |
| | | | Total | | | (8, 8) | t(14)=2.934 | unpaired t-test | ***0.0109*** | * |
| | 4C | Cleaved gephyrin | Synaptic | 7-day Veh vs DZP | | (8, 8) | t(14)=0.1385 | unpaired t-test | 0.8918 | ns |
| | | | Extrasynaptic | | | (8, 8) | - | Mann-Whitney test | ***0.0003*** | *** |
| | | | Total | | | (8, 8) | t(14)=0.2080 | unpaired t-test | 0.8382 | ns |
| | 4D | Cleaved/Full-length gephyrin | Synaptic | 7-day Veh vs DZP | | (8, 8) | t(14)=2.576 | unpaired t-test | ***0.0220*** | * |
| | | | Extrasynaptic | | | (8, 8) | - | Mann-Whitney test | ***0.0003*** | *** |
| | | | Total | | | (8, 8) | t(14)=0.7224 | unpaired t-test | 0.4819 | ns |
| Section 3.4 | 5B | Surface synaptic γ2-GABA$_A$R | Number of Objects | 7-day Veh vs DZP | 3 | (43, 45) | t(86)=2.379 | unpaired t-test | ***0.0196*** | * |
| | | | Binary Area | | | (43, 47) | t(88)=2.073 | unpaired t-test | ***0.0411*** | * |
| | | | Sum Intensity | | | (42, 47) | t(87)=0.8576 | unpaired t-test | 0.3935 | ns |
| | 5C | Surface extrasynaptic γ2-GABA$_A$R | Number of Objects | 7-day Veh vs DZP | | (42, 47) | t(87)=1.412 | unpaired t-test | 0.1616 | ns |
| | | | Binary Area | | | (42, 46) | t(86)=1.864 | unpaired t-test | 0.0657 | ns |
| | | | Sum Intensity | | | (41, 47) | t(86)=2.669 | unpaired t-test | ***0.0091*** | ** |
| | 5D | Total surface γ2-GABA$_A$R | Number of Objects | 7-day Veh vs DZP | | (43, 47) | t(88)=1.047 | unpaired t-test | 0.2981 | ns |
| | | | Binary Area | | | (42, 47) | t(87)=1.143 | unpaired t-test | 0.2561 | ns |
| | | | Sum Intensity | | | (43, 46) | t(87)=0.1481 | unpaired t-test | 0.8826 | ns |
| | 5E | GAD65 | Number of Objects | 7-day Veh vs DZP | | (43, 46) | t(87)=1.575 | unpaired t-test | 0.1189 | ns |
| | | | Binary Area | | | (43, 47) | t(88)=1.028 | unpaired t-test | 0.3068 | ns |
| | | | Sum Intensity | | | (43, 47) | t(88)=0.6840 | unpaired t-test | 0.4957 | ns |
| | 5G | γ2-GABA$_A$R surface and total protein expression by surface biotinylation | Surface | 7-day Veh vs DZP | 6 | (15, 13) | t(26)=1.248 | unpaired t-test | 0.2231 | ns |
| | | | Total | | 9 | (22, 22) | t(42)=0.3200 | unpaired t-test | 0.7505 | ns |
| | S5B | α4-GABA$_A$R expression | Synaptic | 7-day Veh vs DZP | 5 | (10, 10) | t(18)=0.0807 | unpaired t-test | 0.9366 | ns |
| | | | Total | | 3 | (6, 6) | t(10)=2.098 | unpaired t-test | ***0.0623*** | ns |
| Section 3.5 | 6C | Whole-field gephyrin-γ2 PL signal numbers | | 7-day Veh vs DZP | 3 | (47, 37) | - | Mann-Whitney test | ***0.0561*** | ns |
| | 6D | Synaptic γ2-gephyrin PL signal numbers | | | | (46, 39) | - | Mann-Whitney test | 0.3164 | ns |
| | 6E | Extrasynaptic γ2-gephyrin PL signal numbers | | | | (46, 38) | - | Mann-Whitney test | ***0.0270*** | * |
| Section 3.5 | 7B | Synaptic Gephyrin Fluorescence Recovery Intensity | 120 sec | 7-day Veh vs DZP | 3 | Cells: (17, 17) Synapses: (81, 74) | t(153)=0.01795 | unpaired t-tests | 0.9857 | ns |
| | | | 240 sec | | | | t(153)=0.1654 | | 0.8688 | ns |
| | | | 360 sec | | | | t(153)=0.6786 | | 0.4984 | ns |
| | | | 480 sec | | | | t(153)=1.844 | | 0.0671 | ns |
| | | | 600 sec | | | | t(153)=1.331 | | 0.1851 | ns |
| | | | 720 sec | | | | t(153)=2.007 | | ***0.0465*** | * |
| | | | 840 sec | | | | t(153)=2.255 | | ***0.0255*** | * |
| | | | 960 sec | | | | t(153)=1.992 | | ***0.0481*** | * |
| | | | 1080 sec | | | | t(153)=2.093 | | ***0.0380*** | * |
| | | | 1200 sec | | | | t(153)=2.135 | | ***0.0344*** | * |
| | | | 1320 sec | | | | t(153)=2.336 | | ***0.0208*** | * |
| | | | 1440 sec | | | | t(153)=2.37 | | ***0.0190*** | * |
| | | | 1560 sec | | | | t(153)=2.137 | | ***0.0342*** | * |
| | | | 1680 sec | | | | t(153)=1.895 | | 0.0600 | ns |
| | | | 1800 sec | | | | t(153)=2.031 | | ***0.0439*** | * |
| | | Synaptic γ2-GABA$_A$R Fluorescence Recovery Intensity | 120 sec | 7-day Veh vs DZP | 3 | Cells: (17, 17) Synapses: (81, 73) | t(154)=0.5915 | unpaired t-tests | 0.5550 | ns |
| | | | 240 sec | | | | t(154)=0.4519 | | 0.6520 | ns |
| | | | 360 sec | | | | t(154)=0.2598 | | 0.7953 | ns |
| | | | 480 sec | | | | t(154)=0.8749 | | 0.3830 | ns |
| | | | 600 sec | | | | t(154)=0.1169 | | 0.9071 | ns |
| | | | 720 sec | | | | t(154)=0.07697 | | 0.9387 | ns |
| | | | 840 sec | | | | t(154)=0.8724 | | 0.3844 | ns |
| | | | 960 sec | | | | t(154)=0.1433 | | 0.8863 | ns |

| Supplementary Table S1: Summary of Statistics (page 3 of 3) | | | | | | | | | | |
|---|---|---|---|---|---|---|---|---|---|---|
| **Location in Text** | | **Measurement** | | **Comparison** | ***N* Values (Independent Cultures)** | ***n* values (Veh, DZP)** | **Statistic (df)** | **Statistical test** | ***p* value** | |
| **Section** | **Figure** | | | | | | | | | |
| Section 3.5 | 7B (*cont*) | Synaptic $\gamma$2-GABA$_A$R Fluorescence Recovery Intensity | 1080 sec | 7-day Veh vs DZP | 3 | Cells: (17, 17) Synapses: (81, 73) | t(154)=0.6563 | unpaired t-tests | 0.5126 | ns |
| | | | 1200 sec | | | | t(154)=1.05 | | 0.2952 | ns |
| | | | 1320 sec | | | | t(154)=0.9558 | | 0.3407 | ns |
| | | | 1440 sec | | | | t(154)=0.2453 | | 0.8066 | ns |
| | | | 1560 sec | | | | t(154)=0.5545 | | 0.5800 | ns |
| | | | 1680 sec | | | | t(154)=0.4424 | | 0.6589 | ns |
| | | | 1800 sec | | | | t(154)=0.299 | | 0.7653 | ns |
| | 7D | Extrasynaptic Gephyrin Fluorescence Recovery Intensity | 120 sec | 7-day Veh vs DZP | 3 | (17, 16) | t(31)=0.5553 | unpaired t-tests | 0.5827 | ns |
| | | | 240 sec | | | | t(31)=0.05797 | | 0.9541 | ns |
| | | | 360 sec | | | | t(31)=0.4562 | | 0.6514 | ns |
| | | | 480 sec | | | | t(31)=0.6977 | | 0.4906 | ns |
| | | | 600 sec | | | | t(31)=1.24 | | 0.2244 | ns |
| | | | 720 sec | | | | t(31)=1.152 | | 0.2582 | ns |
| | | | 840 sec | | | | t(31)=1.181 | | 0.2465 | ns |
| | | | 960 sec | | | | t(31)=0.7508 | | 0.4584 | ns |
| | | | 1080 sec | | | | t(31)=0.6102 | | 0.5462 | ns |
| | | | 1200 sec | | | | t(31)=0.1191 | | 0.9060 | ns |
| | | | 1320 sec | | | | t(31)=0.07248 | | 0.9427 | ns |
| | | | 1440 sec | | | | t(31)=0.7137 | | 0.4807 | ns |
| | | | 1560 sec | | | | t(31)=0.2604 | | 0.7963 | ns |
| | | | 1680 sec | | | | t(31)=0.05484 | | 0.9566 | ns |
| | | | 1800 sec | | | | t(31)=0.5287 | | 0.6008 | ns |
| | | Extrasynaptic $\gamma$2-GABA$_A$R Fluorescence Recovery Intensity | 120 sec | 7-day Veh vs DZP | 3 | (17, 17) | t(32)=2.172 | unpaired t-tests | ***0.0374*** | * |
| | | | 240 sec | | | | t(32)=1.413 | | 0.1674 | ns |
| | | | 360 sec | | | | t(32)=2.284 | | ***0.0292*** | * |
| | | | 480 sec | | | | t(32)=1.331 | | 0.1925 | ns |
| | | | 600 sec | | | | t(32)=2.951 | | ***0.0059*** | ** |
| | | | 720 sec | | | | t(32)=2.089 | | ***0.0448*** | * |
| | | | 840 sec | | | | t(32)=2.04 | | ***0.0497*** | * |
| | | | 960 sec | | | | t(32)=2.215 | | ***0.0340*** | * |
| | | | 1080 sec | | | | t(32)=2.533 | | ***0.0164*** | * |
| | | | 1200 sec | | | | t(32)=2.664 | | ***0.0120*** | * |
| | | | 1320 sec | | | | t(32)=2.875 | | ***0.0071*** | ** |
| | | | 1440 sec | | | | t(32)=1.305 | | 0.2011 | ns |
| | | | 1560 sec | | | | t(32)=1.11 | | 0.2752 | ns |
| | | | 1680 sec | | | | t(32)=1.923 | | 0.0634 | ns |
| | | | 1800 sec | | | | t(32)=2.515 | | ***0.0173*** | * |
