## Supplemental Materials and Figures for "Chronic benzodiazepine treatment triggers gephyrin scaffold destabilization and GABA_A_R subsynaptic reorganization"

**Supplementary Materials and Methods**

**Materials**

| **Antibody** | **Host Species** | **Company** | **Identifiers** | **Dilution** | **Experiment** | **Figure** |
| --- | --- | --- | --- | --- | --- | --- |
| GABA_A_R α5 | rabbit | Synaptic Systems | Cat# 224 503 RRID:AB_2619944 | 1/1000 | Western Blot | S4 |
| NMDAR GluN2A | rabbit | Phospho-Solutions | Cat# 1500-NR2A RRID:AB_2492170 | 1/1000 | Western Blot | S4 |
| NMDAR GluN2B | mouse | NeuroMab | Cat# N59/36 RRID:AB_2877296 | 1/1000 | Western Blot | S4 |
| PSD95 | mouse | Thermo Fisher | Cat# MA1-046 RRID:AB_2092361 | 1/2000 | Western Blot | S4 |
| Radixin | mouse | Novus | Cat# H00005962-M06 RRID:AB_547562 | 1/500 | Western Blot | S4 |
| EEA1 | mouse | BD Biosciences | Cat# 610457 RRID:AB_397830 | 1/1000 | Western Blot | S4 |
| β-actin | mouse | Sigma-Aldrich | Cat# A1978 RRID:AB_476692 | 1/2000 | Western Blot | S4 |
| GABA_A_R α4 | rabbit | Phospho-Solutions | Cat# 845-GA4C RRID:AB_2492103 | 1/1000 | Western Blot | S5 |

**Syn-PER synaptic protein extraction and western blotting**

Biochemical enrichment of the synaptosomal fraction using Syn-PER Synaptic Protein Extraction Reagent (Thermo Fisher, Cat# 87793) was performed following the manufacturer’s protocol. Briefly, neurons treated with vehicle or DZP for seven days were rinsed twice with ice-cold DPBS and lysed in Syn-PER reagent supplemented with protease and phosphatase inhibitors. Following a low-speed centrifugation (1,200 x*g*, 10 min, 4°C) to remove cell debris, a small portion of the lysate (total protein fraction) was saved for downstream analysis. Samples were then centrifuged at 15,000 x*g* for 20 min at 4°C to generate the synaptosome pellet, which was resuspended in Syn-PER reagent containing protease and phosphatase inhibitors. Protein concentrations for each fraction were determined by BCA Protein Assay (Thermo Fisher). Equal amounts of protein from the synaptic and total fractions were resolved by SDS-PAGE and transferred overnight to supported nitrocellulose membrane (Bio-Rad). Membranes were incubated overnight at 4°C with primary antibodies against the α4-GABA_A_R subunit and GAPDH. HRP-coupled secondary antibodies were applied for one hour at room temperature followed by chemiluminescent visualization. Analysis was performed using the Image Lab 6.0 (Bio-Rad) volume tool with global background subtraction and α4-GABA_A_R immunoreactivities normalized to that of GAPDH loading control. Within each independent culture, measurements were normalized to the vehicle-treated average.

**Supplementary Figures**

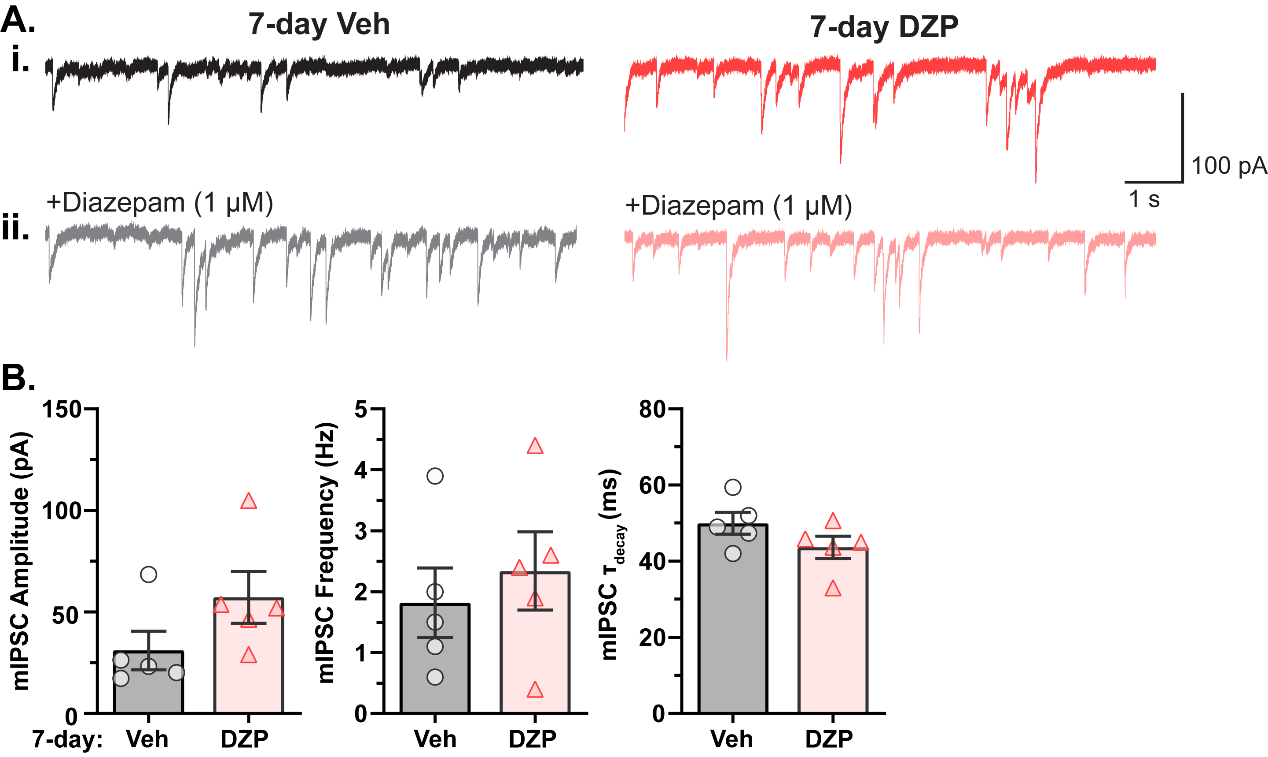

**Figure S1. mIPSC parameters are not altered by chronic DZP treatment in cultured cortical neurons.**

Miniature inhibitory postsynaptic currents (mIPSCs) were measured by whole-cell recordings to assess baseline inhibitory function in neurons treated with vehicle (Veh) or 1 μM DZP for seven days. **(A)** Representative mIPSC traces from 7-day Veh- or DZP-treated cortical neurons i) before and ii) after acute application of 1 μM diazepam. **(B)** Baseline mIPSC amplitude (Veh=31.1±9.47 pA, DZP=57.2±12.7 pA; *p*=0.1379), frequency (Veh=1.8±0.57 Hz, DZP=2.3±0.64 Hz; *p*=0.5615), and τ_decay_ (Veh=50.0±2.87 ms, DZP=43.7±2.92 ms; *p*=0.1618) are similar between 7-day Veh- and DZP-treated neurons.

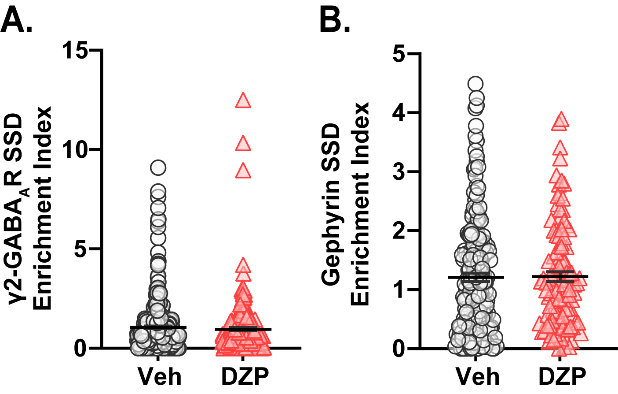

**Figure S2. Chronic DZP treatment does not alter the subsynaptic enrichment indices of γ2-GABA_A_R and gephyrin.**

Enrichment index was calculated as the average local density of one protein within a 60 nm range from an SSD peak of the other protein. γ2-GABA_A_R and gephyrin SSD enrichment indices were similar in 7-day Veh- vs DZP-treated neurons. **(A)** Enrichment of γ2-GABA_A_R SSDs to gephyrin SSDs. **(B)** Enrichment of gephyrin SSDs to γ2-GABA_A_R SSDs. *n*=7-9 cells, N=2 independent cultures; mean ± SEM. Analyses by Mann-Whitney test.

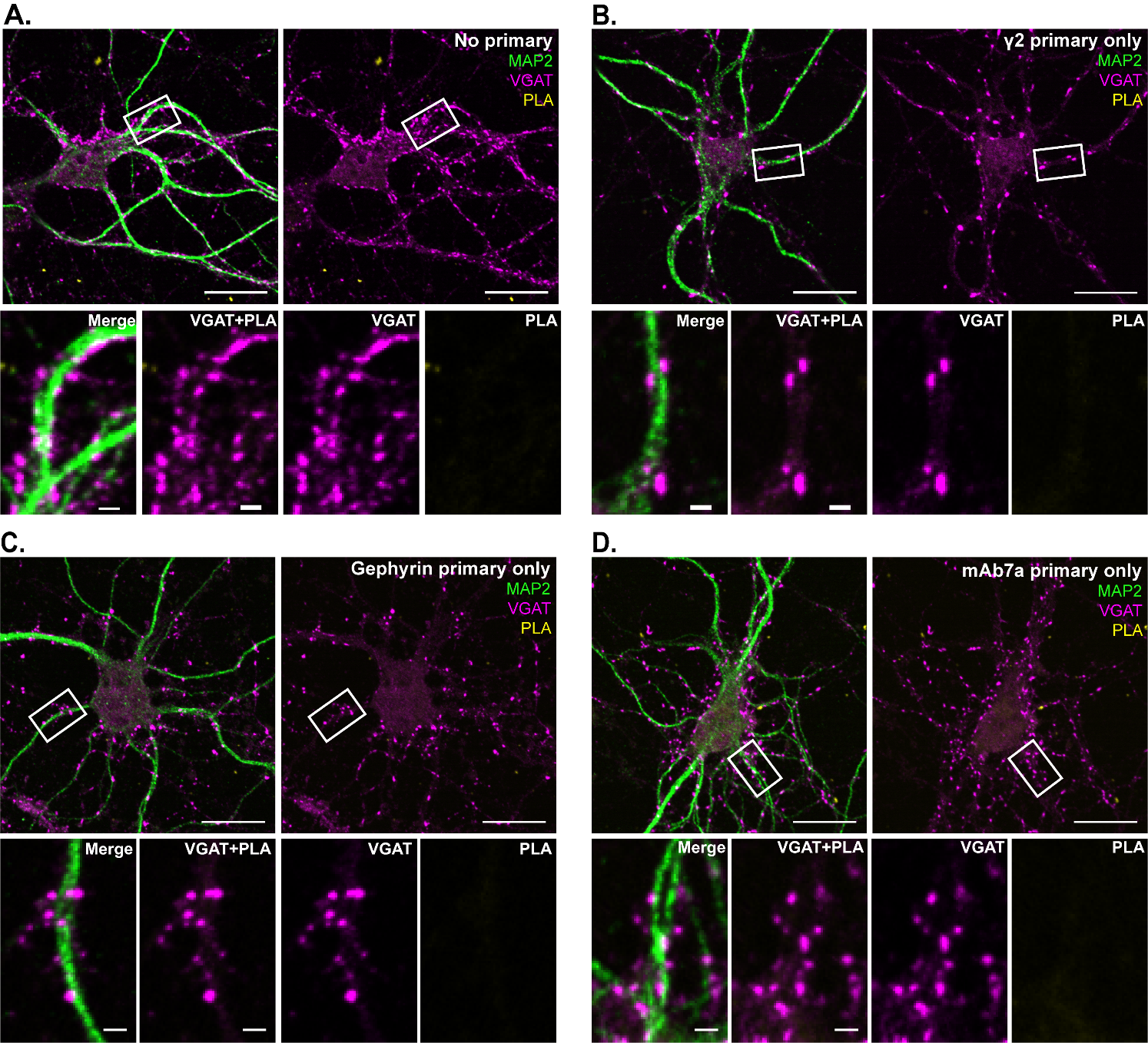

**Figure S3. Minimal PL signal is observed under no-primary or single-primary antibody conditions.**

Following the proximity ligation assay (PLA) protocol, neurons were permeabilized and incubated with either **(A)** no primary antibodies; **(B)** only γ2-GABA_A_R subunit primary antibody; **(C)** only total gephyrin (3B11) primary antibody; or **(D)** only mAb7a primary antibody, followed by incubation with Navenibody secondary reagents. Minimal PL signal (yellow) is observed in each case. Counterstaining was performed with MAP2 (green) and VGAT (pink) to visualize neuronal dendrites and inhibitory synapses, respectively. Scale bars are 20 μm for neurons and 2 μm for dendrite zoom images.

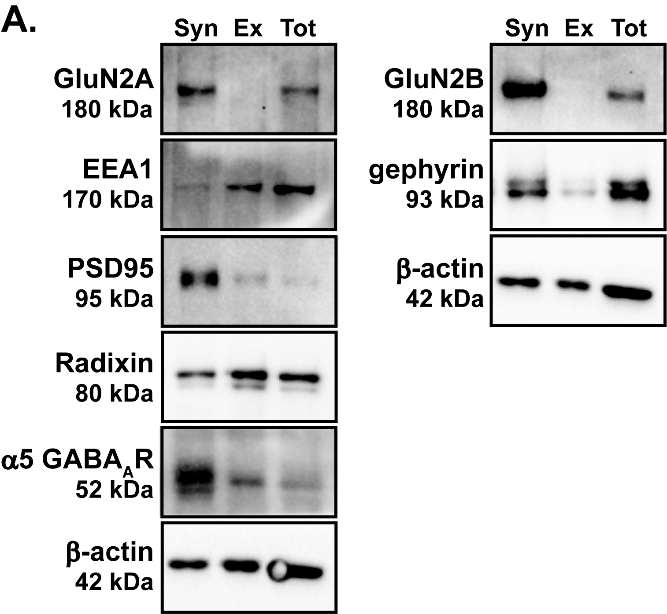

**Figure S4. Validation of the subcellular fractionation protocol.**

To confirm isolation of the synaptic and extrasynaptic membrane fractions using the subcellular fractionation protocol, each fraction was immunoblotted for various synaptic (glutamatergic NMDA receptor subunits GluN2A, GluN2B; excitatory postsynaptic scaffold PSD95; inhibitory postsynaptic scaffold gephyrin) or extrasynaptic (early endosome antigen 1 (EEA1); and radixin, a cytoskeletal protein that clusters extrasynaptic α5-GABA_A_Rs) markers. Syn=synaptic fraction; Ex=extrasynaptic fraction; Tot=total protein lysate.

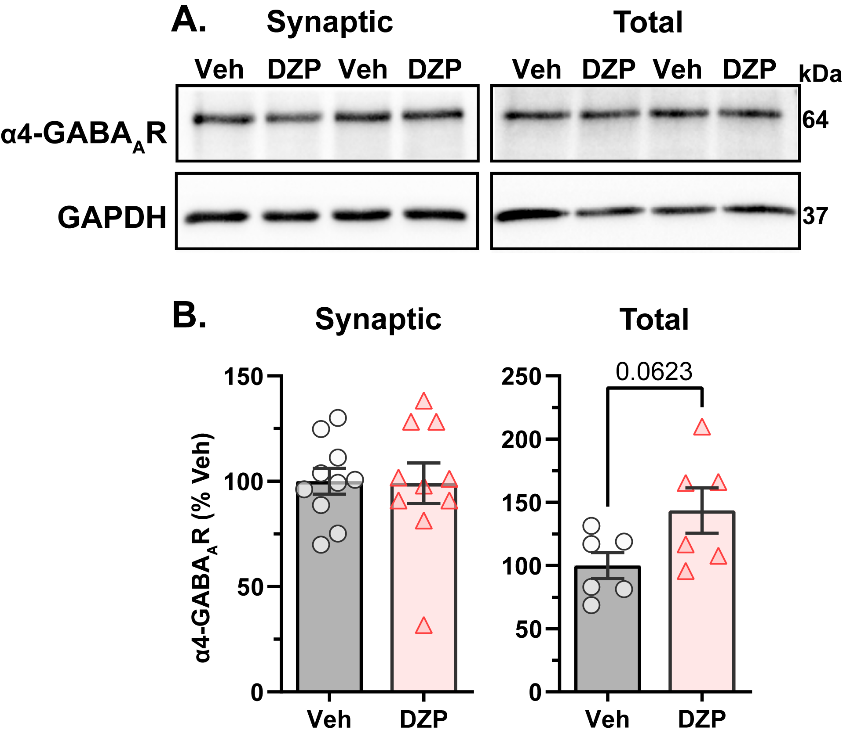

**Figure S5. α4-GABA_A_R synaptic protein expression is unchanged by chronic DZP treatment.**

Synaptic protein extraction (SynPER) and subsequent western blotting was used to assess endogenous expression of the α4-GABA_A_R subunit in the synaptic and total protein fractions of neurons treated for 7 days with Veh or 1 μM DZP. **(A)** Representative SynPER western blots for the synaptic and total fractions. **(B)** Quantification of α4-GABA_A_R immunoreactivities in Veh- vs DZP-treated neurons. Immunoreactivity was quantified with normalization to GAPDH loading control and assessed relative to the vehicle-treated culture average. α4-GABA_A_R expression was not significantly changed in either the synaptic or total fraction after 7-day DZP treatment, though total protein expression trended upward (Veh=100±10%, DZP=144±18%, *p*=0.0623). *n*=2 replicates per culture, N=3-5 independent cultures; mean ± SEM. Analysis by unpaired *t*-test.

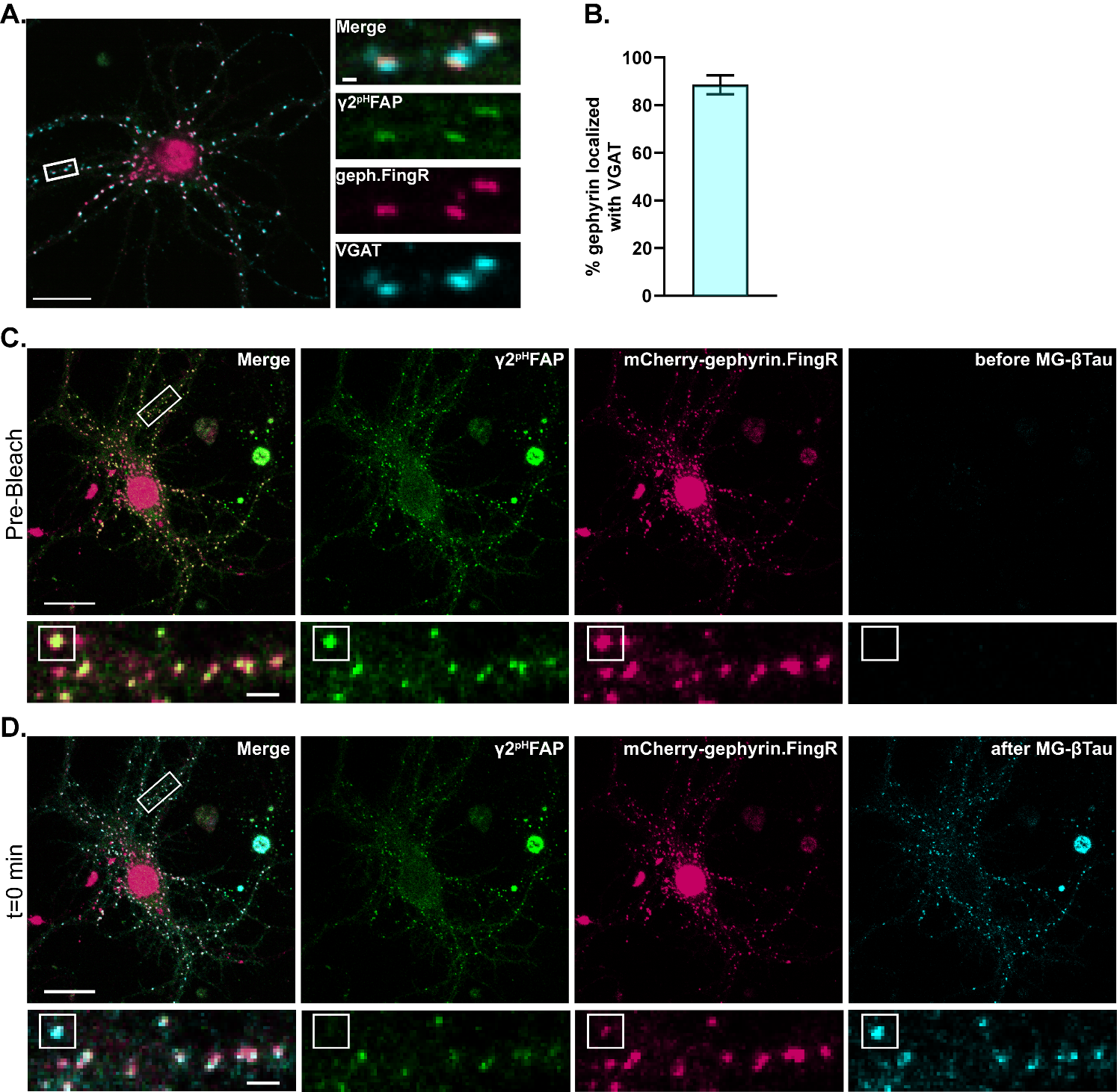

**Figure S6. Confirmation of synaptic localization of mScarlet-Gephyrin.FingR and surface expression of γ2^pH^FAP clusters for FRAP experiments.**

Fluorescence Recovery After Photobleaching (FRAP) experiments were performed in hippocampal neurons co-transfected with γ2^pH^FAP and mScarlet-Gephyrin.FingR (geph.FingR) and treated with Veh or 1 μM DZP for seven days. **(A)** Representative neuron co-transfected with γ2^pH^FAP and mScarlet-Gephyrin.FingR and live-labeled with VGAT CypHer5E. **(B)** Quantification of mScarlet-Gephyrin.FingR colocalized with VGAT CypHer5E confirms predominantly synaptic localization of bright geph.FingR clusters (~90%). **(C,D)** MG-βTau, a cell-impermeant dye that binds to FAP tags with high affinity, was added to neurons immediately after photobleaching to confirm surface expression of γ2^pH^FAP clusters. **(C)** Whole cell and dendrite zoom images before photobleaching and before addition of MG-βTau. **(D)** Whole cell and dendrite zoom images immediately after photobleaching (t=0 min) and after addition of 10 nM MG-βTau. The white box indicates a synaptic cluster that was photobleached and used in analysis; MG-βTau labeling of this cluster confirms γ2^pH^FAP surface localization. A,C,D: Scale bars are 20 μm for neurons and 1 μm (A) or 2 μm (C,D) for dendrite zoom images. B: *n*=24 dendritic segments of 10 μm from 8 neurons, N=1 culture; mean ± SEM.
